# Tumour neoantigen repertoire computational predictions in malignant peripheral nerve sheath tumours define potential targets for immunotherapy

**DOI:** 10.64898/2026.04.05.713607

**Authors:** Mirvat Surakhy, Joseph E. Caesar, Masooma Rajput, Qinan Qian, A. Bassim Hassan

**Affiliations:** Oxford Molecular Pathology Institute, Sir William Dunn School of Pathology, University of Oxford, South Parks Road, Oxford, OX1 3RE, United Kingdom

**Keywords:** Tumour antigen, Genomics, Malignant Peripheral Nerve Sheath Tumour, MPNST, Neurofibromatosis, NF1, persistent tumour mutational burden, pTMB, Major histocompatibility Complex, MHC, T-cell receptor, TCR, immune therapy, tumour microenvironment

## Abstract

Malignant peripheral nerve sheath tumours (MPNSTs) are high grade soft-tissue sarcomas with an unmet need for novel therapies. Tumour antigen-based approaches, including neoantigen and tumour-associated antigens (TAA) directed therapies, offer potential opportunities for immunotherapy. Here, we integrated public domain tumour DNA and RNA sequencing data with in-silico predictions in order to characterise the potential (neo)antigenic landscape of MPNST. We stratified the computational predictions across the two known sub-groups of MPNST, those associated without and with Polycomb Repressor Complex 2 (PRC2) loss of function variants (PRC2-Loss). Using pVACtools, computationally identified high-confidence neoantigens based on predicted pMHC affinity were derived from somatic mutations and gene fusions, as well as recurrently overexpressed cell-surface TAAs. All predicted neoantigens were private to individual MPNST cases and across both tumour subtypes. Using ImSig and CIBERSORTx, PRC2-Loss tumours displayed reduced immune infiltration with downregulation of antigen processing and presentation pathways compared to PRC2-WT, confirming known intrinsic constraints to effective neoantigen-directed immune priming. Moreover, PRC2-Loss MPNSTs demonstrated recurrent copy number driven overexpression of cell surface TAAs derived from chromosome 8 amplification, providing potential immunotherapeutic targets that are pMHC independent. Overall, these predictive findings confirm a PRC2-independent private immuno-antigenic peptide repertoire, with an immune resistant MPNST microenvironment in PRC-Loss. These data provide further impetus for personalised functional validation and immune based treatment strategies, including personalised neoantigen vaccines and cell surface protein TAA-directed therapies dependent on PRC2 status.

## Introduction

Malignant peripheral nerve sheath tumours (MPNSTs) are highly aggressive soft tissue sarcomas derived from either Schwann cells or pluripotent cells of the neural crest (Suppiah et al., 2023). Complete surgical excision of MPNSTs remains the only proven curative treatment when achievable. Clinical trials using therapies targeting dysregulated pathways, such as epidermal growth factor receptor (EGFR) inhibitors, angiogenesis inhibitors, MAPK/MEK pathway and mTOR inhibitors, have resulted in marginal and non-sustainable clinical benefits, underlining the unmet need for new therapeutic options (Prudner et al., 2020).

MPNSTs mainly occur either sporadically or in the context of neurofibromatosis type 1 (*NF1*, 5-15%), an autosomal dominant tumour predisposition syndrome where lethality is often associated with subsequent MPNST development (Evans et al., 2002). Mutations of *NF1* activate the RAS pathway leading to an increase in the activity of PI3K/AKT/mTOR and/or MEK-ERK which drive cell proliferation and survival (Baez-Flores et al., 2023) (Gonzalez-Munoz et al., 2022). Bi-allelic loss of function of *NF1*, resulting in loss of the RAS-Gap protein neurofibromin (NF1), is frequently followed by loss of function of Cyclin Dependent Kinase 2A/2B (*CDKN2A*), components of the Polycomb Repressive Complex 2 (PRC2) such as either Embryonic Ectoderm Development (*EED*) or Suppressor of Zeste 12 homolog (*SUZ12*) and the tumour suppressor Tp53 (*TP53*). Together, these gene pathways are considered the main drivers of MPNST (Cortes-Ciriano et al., 2023; Lee et al., 2014; Magallon-Lorenz et al., 2023; Yan et al., 2022). Importantly, the loss of histone H3 lysine 27 trimethylation (H3K27me3) mediated repressive chromatin occurs in approximately 50% of MPNSTs and stratifies MPNSTs into two potential mechanistic groups (Cortes-Ciriano et al., 2023; Lee et al., 2014; Yan et al., 2022). PRC2 complex mutations in either *EED* or *SUZ12* are mutually exclusive, and result in reduction in Enhancer of Zeste Homolog1/2 (*EZH1/2*) subunit mediating methyl transferase activity (H3K27me3). The resulting widespread alterations in gene expression also include regulatory genes for immune cell infiltration, genomic instability and the selection for subsequent amplification of chromosome 8 (Cortes-Ciriano et al., 2023; Yan et al., 2022; Zhang et al., 2024). Moreover, MPNSTs associated with a wild-type PRC2 complex appear to activate the RAS-MEK-ERK, WNT/β-catenin/CCND1 pathways, whereas MPNSTs with loss of function of PRC2 appearing to activate the sonic hedgehog (SHH) pathway (Suppiah et al., 2023). Importantly, the stratification of MPNSTs by PRC2 loss of function status (PRC2-Loss versus PRC2-WT) by either loss of H3K27me3 immuno-detection or by presence of PRC2 gene variant sequencing, suggests that PRC2-Loss is predicted to be associated reduced cytokine gene expression, reduced immune cell infiltration and T-cell priming (Yan et al., 2022).

Cancer neoantigens can result from translation of non-synonymous single nucleotide, translocation breakpoint and alternative splicing coding and non-coding variants (Xie et al., 2023). The intra-cellular peptides generated and that are proteolytically processed can be presented at the cell surface of antigen presenting cells by major histocompatibility (MHC) class I and class II molecules. T-cell receptor recognition of the presented neoantigen peptide-MHC (pMHC) complex can initiate an adaptive immune response against cancer cells (Xie et al., 2023). Not surprisingly, avoiding immune recognition and destruction is one of the hallmarks of cancers including MPNSTs (Dagogo-Jack and Shaw, 2018; Hanahan, 2022; Hanahan and Weinberg, 2011). Mechanisms include escape from T-cell killing by upregulating negative checkpoint regulators of the TCR response, such as programmed cell death ligand -1 (PDL-1) that binds to PD-1 and is predominantly expressed on the surface of antigen-stimulated T cells (Yamaguchi et al., 2022). Tumour cells can also evade T-cell-mediated cytotoxicity by either downregulating the expression of MHC-I/II and mutating or deleting the HLA locus required for pMHC presentation (Cornel et al., 2020). Immune checkpoint blockade (ICB) using anti-PDL-1/anti-PD-1 alone, or in combination with anti-cytotoxic T-lymphocyte-associated protein 4 (CTLA4), can reinstate T cell cytotoxicity resulting in sustained disease control in melanoma, lung and other solid tumours. Moreover, tumour mutation burden (TMB) is a correlative biomarker for predicting patient responses to ICB in some tumour types (Chan et al., 2019; Hiam-Galvez et al., 2021; Rotte, 2019), but is not raised significantly in MPNSTs and the responses to ICB therapy are difficult to predict (Larson et al., 2022).

Next generation sequencing (NGS) of MPNSTs (DNA whole genome, whole-exome and RNA-Seq) have recently characterised the genomes of these high-grade tumours, yet the potential neoantigens that may be presented to the host adaptive immune system have yet to be fully characterised (Cortes-Ciriano et al., 2023). Comprehensive NGS of tumour-normal pairs combined with mRNA expression have greatly facilitated the identification of expressed patient specific-neoantigen repertoires when integrated with machine learning prediction algorithms. Computational prediction of peptide binding affinity to different MHC molecules expressed from patient specific HLA alleles and are based on models derived from large structural and immuno-peptidomic data sets. Such predictive approaches have led to selection of neoantigens with predicted high-binding affinity to MHC molecules, but are unable to predict the TCR-variable chain sequences that bind the pMHC, yet are used as the basis for personalised cancer mRNA vaccines (Lang et al., 2022). Subsequent validation of polyclonal TCR responses in this clinical context estimates neoantigen predictions of significance in approximately 25% of cases. Specific neoantigen-based vaccines also prove to be more immunogenic overall and not affected by central immunological tolerance compared to vaccines targeted towards tumour-associated antigens (TAA), where self-antigens are only differentially (highly) expressed by tumours compared to normal cells (Melero et al., 2014; Peng et al., 2019). While neoantigen presentation depends on HLA expression and MHC presentation, TAAs can also be clinically relevant MHC independent immunotherapeutic targets. For instance, chimeric antigen receptor (CAR) T-cells targeting cell surface antigens have demonstrated growing promise for solid tumours treatment by enabling T-cell recognition of surface antigens in a manner that is independent of MHC presentation. CARs rely on antibody-derived binding domains, and their activity depends critically on identifying TAAs that are highly expressed on tumour cells while exhibiting minimal expression in normal tissues (Kandra et al., 2022).

Despite such immunotherapeutic approaches, a comprehensive characterisation of predicted MPNST associated neoantigen presentation has yet to be completed. There are essentially two complimentary approaches for neoantigen identification that are greatly dependent on sample availability and processing. The first is an MHC dependent immuno-peptidomic analysis using mass spectrometry, where ideally large amounts of fresh tumour sample availability circumvents the challenge of detection of rarer peptide-MHC complexes (Bassani-Sternberg, 2018; Shapiro and Bassani-Sternberg, 2023; Shapiro et al., 2025). The second approach, where there are no tumour samples available, is in silico computational approach based on RNA expression, that has increasingly been adopted because of particular and limited sample availability in rarer cancers, such as MPNSTs. Here, we report our initial computational prediction analysis using two sequencing cohorts of MPNST tumours. This approach based on a series of machine learning algorithms trained on pMHC structure, peptidomic and functional data, where affinity predictions are restricted to expressed MHCs (Okada et al., 2022). We report our analysis genomic DNA and mRNA MPNST sequencing data using predictive algorithm pipeline, PVACtools. By pooling two MPNST NGS cohorts, the international Genomics of MPNST (GeM) consortium and the database of Genotype and Phenotype (dbGaP), we report the prediction of public and private neoantigen variants and TAA as candidate immunological targets that may be the basis of future MPNST immunotherapy approaches.

## Methods

### Study cohorts

Two independent NGS cohorts of malignant peripheral nerve sheath tumours (MPNST) were evaluated. The first cohort was from dbGaP (phs000792/GRU-PUB-MDS, request number #101010-2) comprised of 16 patients. Samples were of 12 fresh frozen tumours with and adjacent normal tissue from 10 patients subject to whole exome sequencing (WES) and RNA-seq (Lee et al., 2014). The sequencing data was provided in Fastq format. The exome was captured using Agilent SureSelect V4+UTRs exome bait (Agilent), and sequencing was performed on the Illumina HiSeq-2500 platform with 75bp paired-end (PE) reads. The RNA was poly-A selected and processed using the TruSeq RNA sample preparation kit and sequenced on an Illumina HiSeq-2500 platform with 51 bp PE reads. For whole exome sequencing, data quality control was checked using fastqc https://www.bioinformatics.babraham.ac.uk/projects/fastqc/ (v 0.11.9). Adapters were marked using MarkIlluminaAdapters (Picard) wrapper implemented in the genome analysis tool kit (GATK, v 4.2.3.0). Sequencing reads were mapped to the GRCh38 human reference genome using BWA-MEM (v 0.7.17) (Li and Durbin, 2009) followed by remove duplicates and base quality score recalibration following the GATK best practices workflow (McKenna et al., 2010). Alignment and coverage metrics as well as PCR duplicate marking were computed using Picard tools (v1.125).

The second larger cohort was from the international Genomics of MPNST (GeM) consortium, the data was available via the European Genome-phenome Archive (EGA) with accession number EGAD00001008608) (Cortes-Ciriano et al., 2023). The data was from 88 patients. Seventy-six patients had fresh frozen tumours and adjacent normal tissue with whole genome sequencing (WGS) and RNA-seq. DNA sequencing libraries were prepared using the TruSeq DNA PCR Free 350bp kit, which was sequenced on the Illumina NovaSeq6000 sequencing machine to generate 2x 151 PE reads. The provided files were in .bam format that mapped to the GRCh38 build human reference genome and processed following the Genome Analysis Toolkit (GATK, version 4.1.8.0) best practices workflow to remove duplicates and recalibrate base quality scores (McKenna et al., 2010). RNA was poly-A selected and libraries were prepared using the Illumina TruSeq Strand Specific Large Insert RNA kit (50 M pairs) v1. Libraries were sequenced on the Illumina HiSeq4000 sequencing machine at the Broad Institute (Cambridge, MA) with 101 bp PE reads. RNA-seq files provided in Fastq format. Throughout the manuscript and figures, ‘cases’ represent unique patients and may comprise one or more tumour samples; sample*s* refer to individual separate tumour specimens used for sequencing analysis.

### Germline SNVs and INDELs

The germline short variant discovery workflow from GATK (v4.2.3.0) was used to identify germline SNVs and INDELs from the normal samples (WGS/WES). The intermediate GVCF files were generated for each sample using HaplotypeCaller in GVCF mode. For the WES, the interval list for the SureSelectV4+UTRs S03723424 kit lifted to the hg38 reference genome with100bp padded interval. Next, GVCF files for all samples were consolidated into a single GVCF file using the CombineGVCFs with default options. Finally, all samples were jointly genotyped using GenotypeGVCFs using default options. Hard filtering was applied for the dbGaP cohort (for SNPs: --filter-expression QualByDepth (QD)<2.0, FisherStrand (FS)>60.0, RMSMappingQuality (MQ)<40.0, MappingQualityRankSumTest (MQRankSum)<-12.5, ReadPosRankSumTest (ReadPosRankSum)<-8.0, StrandOddsRatio (SOR)>3.0. For Indels --filter-expression QD<2.0, FS >200.0, ReadPosRankSum <-20.0, SOR >10.0). For WGS, VariantRecalibration on SNPs and Indels was performed. The SNP recalibration model was built using the hapmap_3.3.hg38.vcf, 1000G_omni2.5.hg38.vcf, 1000G_phase1snps.high_confidence. hg38.vcf, and Homo_sapiens_assembly38.dbsnp138.vcf datasets and the Indels recalibration model using the Mills_and_1000G_gold_standard.indels.hg38.vcf and the Homo_sapiens_assembly38.dbsnp138.vcf according to the GATK pipeline. ApplyVQSR for SNPs and Indels was performed using the truth-sensitivity-filter-level of 99.5. Ensembl Variant Effect Predictor (VEP) (version 105) was used to annotate all variants.

### Somatic SNV and INDEL detection

Sample pre-processing and mapping were performed as described before. Then somatic SNVs and indels were called using Strelka2 (v2.9.10) (Kim et al., 2018), Varscan2 (v2.4.2) (Koboldt et al., 2012), Mutect2 (Benjamin D et al., 2019). These variant callers were run with default parameters. For the WES, the interval list is described above, and for the WGS, we restricted our analysis to chr1:22, Y, X and M and excluded the black list genes (downloaded from https://dozmorovlab.github.io/excluderanges/#bedbase-data-download). The variants were then combined using the GATK’s CombineVariants and normalised using GATK’s LeftAlignAndTrimVariants to left-align the indels and trim common bases. Then, Vt (Tan et al., 2015) was used to split multiallelic variants, and the GATK’s SelectVariants to select the PASS variants. The VCF files were annotated using VEP (parameters: --format VCF --plugin Downstream --plugin Frameshift --plugin Wildtype --symbol --term SO --transcript_version --tsl --coding_only --pick --hgvs) following the recommendation from PVACtools. Variant coverage for both the tumour and normal DNA data was evaluated using bam-readcount (V1.1.1) https://hub.docker.com/r/mgibio/bam_readcount_helper-cwl/tags (Khanna et al., 2021) for SNPs and indels separately. Then, the data from the output files were added to the annotated VCF using vcf-readcount-annotator from VAtools (v5.0.1) (http://vatools.org). Phased-VCF files were generated by combining both the somatic and germline variants and running the sorted combined variants using the GATK’s ReadBackedPhasing (GATK v3.6.0) as recommended by PVACtools.

### Structural variation analysis

Structural variation was analysed from WGS using Manta (v1.6.0; Python-2.7.1) from paired tumour and normal DNA samples, following the recommended settings. The variants were then annotated using VEP (v105.0) with the –plugin StructuralVariantOverlap option and nstd166.GRCh38.variant_call.vcf.gz reference dataset.

### RNA-seq analysis

Following data QC using fastqc we aligned the patient’s tumour RNA-seq data using HISAT2 (Kim et al., 2019). Stringtie (Pertea et al., 2015) and Kallisto (Bray et al., 2016) were employed for evaluating both the gene and transcript expression values, respectively. Moreover, coverage support for variants in RNA-seq data was determined with bam-readcount (V1.1.1) and gene and transcript levels were added to the VCF using vcf-expression-annotator from VAtools (v5.0.1).

### Fusion gene calling and annotation

Fusion transcript identification was performed using STAR-Fusion (v1.10.0) (Haas et al., 2019). The CTAT Genome Lib (GRCh38_gencode_v37_CTAT_lib_Mar01202.plug-n-play) was downloaded on Jan 2022 (https://data.broadinstituteorg/Trinity/CTAT_RESOURCE_LIB/). PE fastq files mapped with the CTAT Genome Lib (parameters: --FusionInspector inspect --examine_coding_effect). Fusion pairs irrelevant to cancer and could be false positives have been removed. These included genes annotated with GTEx_recurrent_STARF2019, HGNC_GENEFAM, DGD_PARALOGS, Greger _Normal, Babiceanu_Normal, BodyMap, and ConjoinG databases. We also excluded fusion genes that identified in the normal nerve. Fusion genes (star.fusion.abridge) were annotated using Annotate Gene Fusion (AGFusion, v1.252) and Ensembl reference genome database; homo_sapiens.87.db (parameters: -- middlestar to indicate the fusion position in the fusion peptide sequence. -noncanonical to include information from all possible transcripts).

### HLA typing

RNA-seq Fastq files were used for *in silico* HLA typing for MPNST patients of both cohorts. We used OptiType (v1.3.5) (Szolek et al., 2014) and HLA-HD (v1.4.0) (Kawaguchi et al., 2017) for class-I and class-II HLA typing, respectively, using the default parameters.

### Neoantigen identification using pVACtools

We used pVACseq (v4.0.6) from pVACtools (Hundal et al., 2020) to identify and prioritise neoantigens from somatic missense, in-frame insertion, in-frame deletion, and frameshift variants from a VEP-annotated tumour VCFs files using the default parameters unless specified. Briefly, we used MHCflurry MHCflurryEL MHCnuggetsI MHCnuggetsII NNalign NetMHCIIpan NetMHCIIpanEL NetMHCcons NetMHCpanEL SMM SMMPMBEC and SMMalig epitope prediction algorithms. We used NetMHC NetMHCpan PickPocket instead of their consensus method NetMHCcons for samples that failed with this error message (netmhccons_1_1_python_interface.py:86). We used the median MT score and the median fold change across all chosen prediction methods as the main scoring metric (--top-score-metric=median), and processed VCF entries with a PASS status (--pass-only). Class I sub-peptides were examined across 8-11 k-mers and class II across 12-22 k-mers. We enabled reference-proteome similarity analysis to remove peptides that match normal proteins (--run-reference-proteome-similarity) and provided a reference peptide FASTA for matching-Homo_sapiens.GRCh38.pep.all.fa.gz. The pipeline used the default binding threshold of IC_50_ <500nM (--binding-threshold 500) and applied the standard pVACseq filtering pipeline in order: Binding Filter, Coverage Filter, Transcript Filter, Top Score Filter. For tiering and final filtered output, the best peptide met these quality criteria: gene/transcript expression ≥1, transcript support level (TSL)=1 clonal DNA VAF ≥0.20 (--tdna-vaf 0.20), RNA depth ≥10 (--trna-cov 10) and RNA VAF cutoff used for allele expression calculations (--trna-vaf 0.25). Peptides with a match to the reference proteome (RefMatch=True) were flagged and excluded from the final output file, TUMOUR.filtered.tsv.

For predicting the neoantigens resulting from gene fusions we used pVACfuse from pVACtools (v4.0.5) using the default parameters to process the AgFusion annotated fusion genes. We used the epitope prediction algorithms mentioned earlier for PVACseq prediction. Predicted peptides with binding (IC_50_<500 nM) were considered strong binders. We enabled reference-proteome similarity analysis to remove peptides that match normal proteins (--run-reference-proteome-similarity). Following the run, predicted fusion genes that were not relevant to cancer were removed during filtering steps from TUMOUR.filtered.tsv output file. Specifically, we excluded (genes/peptides) derived from fusion genes annotated in non-cancer or normal tissue datasets, including, GTEx_recurrent_STARF2019, HGNC_GENEFAM, DGD_PARALOGS, Greger _Normal, Babiceanu_Normal, BodyMap, and ConjoinG databases. In addition, we excluded fusion genes that identified in the normal nerve. Moreover, all predicted neoantigens derived from fusions that did not pass FusionInspector validation were excluded from subsequent analyses to ensure the robustness and specificity of fusion-derived neoantigen detection.

### Copy number analysis

Copy number alterations were analysed from WGS and WES using the CNVkit (Talevich et al., 2016) v(0.9.10) following the recommended setting. Briefly, for the WGS, a reference was created from 30 normal tissue sequencing files, which was then used to call CNV from all tumours using cnvkit.py batch -m wgs (parameters: bin size 1000bp, --drop-low-coverage, and ignoring the black list region of the genome). The black list was downloaded from https://xfer.genome.wustl.edu/gxfer1/project/cancer-genomics/copyCat/GRCh38/gaps.bed. We used tumour purity reported previously for the GeM dataset (cd-22-0786_supplementary_tables_s1_and_s2_suppst1.xlsx) (Cortes-Ciriano et al., 2023). For samples without purity data, we used Sequenza (Favero et al., 2015) to determine it.

Similarly, CNV from WES was called using the following parameters (cnvkit.py batch -m hybrid, all normal tissue sequencing files used to build the reference and target regions was based on SureSelectV4UTRs kit with 50bp padded to each side). We determined the purity using Sequenza (Favero et al., 2015).

In both cohorts, absolute copy number were determined using CNVkit’s call command in clonal mode, incorporating sample purity values. For genes overlapping more than one copy number segment, the mean copy number was calculated using the findOverlaps function from the GenomicRanges R package (v1.50.2). Heatmaps of copy number data were generated using CNVkit’s heatmap function applied to .cnr files.

### Patient stratification

Mutations (SNV and SV) and total copy number for *SUZ12, EED, EZH2, EZH1* were retrieved from cd-22-0786_supplementary_tables_s1_and_s2_suppst1.xlsx (Cortes-Ciriano et al., 2023). If there was a pathogenic mutation in *SUZ12, EED, EZH2* and *EZH1* with LOH and/or the total copy number of SUZ12, EED or EZH2 was 0, the sample was labelled as PRC2-loss, otherwise the sample was labelled PRC2 Wild Type (WT). Mutations were verified from our call sets and in case of discrepancies we used our data. Samples that were not classified in the table were identified from our call sets. RNA samples lacking matched tumour DNA were classified according to the PRC2 status of their corresponding parent case. The PRC2 status for the dbGaP cohort was derived from the original publication (Lee et al., 2014) and was based on targeted sequencing in addition to WES.

### Copy number Aggregation and Classification

Gene-level copy number estimates were aggregated at the patient level. For patients with more than one sequenced tumour sample, copy number values were collapsed by calculating the mean copy number for each gene across all available samples. These patient-level mean values were subsequently classified into three copy number states: loss (cn ≤ 1), neutral (cn=2), or gain (cn ≥ 3). All downstream analyses were performed using these aggregated, patient-level copy number profiles to avoid pseudo-replication from multiple samples per individual.

### Identification of overexpression of cell surface antigens

Copy number data from the GeM and dbGaP cohorts were combined and integrated with matched tumour RNA-seq expression data. Rather than performing an unbiased genome-wide antigen discovery, we employed a hypothesis-driven prioritisation strategy focusing on chromosome 8, a region recurrently amplified in MPNST, to identify dosage-driven, tumour-intrinsic surface antigens with therapeutic relevance. Candidate genes were restricted to those encoded on chromosome 8 that exhibited copy number amplification and were annotated as cell-surface proteins using a curated human surfaceome generated through a machine-learning-guided proteomics approach from the Wollscheid laboratory Chromosome 8 subset (3591 genes) (Bausch-Fluck et al., 2018) (data accessed November 2022).

Where multiple tumour samples were available for a given patient, copy number and RNA expression values were aggregated using the mean value. Normality of RNA expression values was assessed using the Shapiro-Wilk test, which indicated that expression distributions deviated from normality. Thus, non-parametric statistical tests were applied throughout. Associations between copy number and RNA expression were assessed using Spearman rank correlation. Resulting p-values were adjusted for multiple testing using the Benjamini-Hochberg false discovery rate (FDR), with statistical significance defined as FDR <0.05; correlations with rho ≥0.5 were considered moderate to strong.

### Immune Cell profiling

Gene-level expression data were used to infer relative immune-cell abundance using ImSig (v1.1.3). The presence and co-expression of ImSig signature genes were first assessed using the gene_stat function with the default correlation threshold (r≥0.6), followed by feature selection using the feature_select function. Immune cell abundance was estimated using the imsig function, generating relative scores for seven immune cell types and three biological processes per sample. Statistical differences in ImSig scores between groups were assessed at the case level using the Wilcoxon rank-sum test (also known as Mann-Whitney U) with FDR adjustment.

### Immune Cell Deconvolution Using CIBERSORTx

Transcript abundances of dbGAP and GEM bulk-RNA seq samples were quantified using Salmon (v1.10.3, GCC-12.3.0 build) in quasi-mapping mode with selective alignment (Patro et al., 2017). Sequence-specific bias and GC bias correction were enabled. The library type was automatically inferred. Gene-level expression matrices were subsequently generated in R (v4.4.1) for downstream analyses (Team, 2024).

Immune cell composition was estimated using CIBERSORTx with the LM22 leukocyte gene signature matrix, which defines 22 human immune cell subsets (Chen et al., 2018). Gene expression matrix was uploaded to the CIBERSORTx web portal (https://cibersortx.stanford.edu/), and deconvolution was performed using support vector regression with 1,000 permutations to assess statistical confidence. Absolute mode was enabled to generate absolute immune scores, allowing comparison of total immune infiltration across samples independent of relative proportions. Quantile normalization was disabled as recommended for RNA-seq input data. All remaining parameters were set to default. Downstream analysis and processing of CIBERSORTx output were performed using R.

### Statistical analysis for gene expression

Group-wise comparisons of gene expression were performed using non-parametric tests on patient-level mean expression values. Differences across three copy number states (gain, neutral, and loss) or MPNST subgroups were assessed using the Kruskal-Wallis test. For genes with significant overall effects (p<0.05), post-hoc Dunn’s tests were conducted with Holm adjustment for multiple testing and considered significant if adjusted p (padj)<0.05. For two-group comparisons, the Wilcoxon rank-sum test was used. Where analyses were restricted to a limited set of pre-specified candidate genes, unadjusted p-values were reported in accordance with established statistical guidance (Rothman, 1990). Fold differences were calculated using the median expression of each group.

### Survival Analysis

Overall survival (OS) was analysed using survival objects constructed from time-to-event and censoring information. Neoantigens predicted by pVACseq and pVACfuse were summed at the sample level and averaged across samples to derive a patient-level neoantigen burden. Patients were stratified into three neoantigen burden groups: low (≤5), medium (6-15), and high (≥16). Kaplan-Meier survival curves were estimated using the survfit function, and differences in OS between groups were assessed using the Mantel-Haenszel log-rank test. All analyses were performed in R (v4.2.3) using the survival (v3.5-3) and survminer (v0.5.0) packages.

### Sample matching validation

As a quality control, we used Sample matching using SNPs in humans (SMaSH) (Westphal et al., 2019) implemented in maftools (Mayakonda et al., 2018) to confirm samples are related. Trio samples (tumour/normal DNA and RNAseq) required for neoantigen discovery using pVACseq, and for the TAA discovery. Cases that did not show correlation in one of the three sample trios were excluded from the study. RNA-seq samples that did not show correlation of being from the same case are also excluded from comparison for pvacfuse.

## Results

### Study cohort and inclusion criteria

Two independent existing NGS datasets were combined for in silico analysis, the dbGaP dataset comprising 16 cases sequenced by WES, and the GeM dataset comprising 88 cases sequenced by WGS. The dbGaP cohort included 15 matched tumour-normal exome pairs and 16 tumour RNA-seq samples. The GeM cohort included 86 cases with matched normal DNA, 105 tumour DNA samples, and 132 tumour RNA-seq, reflecting multi-region and replicate sampling (Supplementary Figure 1a). As neoantigen and TAA discovery requires integration of multiple sequencing modalities, we applied stringent quality control and inclusion criteria across both cohorts. Sequencing quality was assessed using FastQC, and sample identity and relatedness were validated across WES and WGS data using a panel of 6,059 exonic SNPs (SMaSH) (Westphal et al., 2019) (Supplementary Figure. 1b). Samples failing SMaSH identity or relatedness checks were excluded (11 cases corresponding to 14 samples). Additional samples were excluded if tumour RNA and/or matched tumour-normal DNA required for downstream analyses were unavailable (14 cases, 16 samples). The final numbers of excluded and included samples for each analysis pipeline are summarised in Supplementary Figure 1 and Supplementary Table 1. While previous studies stratified MPNSTs based on H3K27me3 loss or retention inferred from methylation profiling (Cortes-Ciriano et al., 2023), we directly classified tumours using genomic variant alterations affecting core PRC2 gene components (e.g. *SUZ12, EED, EZH1/2*).

### Patient-Specific Somatic-Neoantigen Landscape and HLA Diversity

Coding somatic mutations from missense variants, in-frame insertions and deletions, protein-altering substitutions and frameshifts, represent the major mRNA coding source of tumour neoantigens. We characterised the neoantigenic epitope repertoire of MPNST tumours arising from these variants using pVACseq (personalised Variant Antigens by Cancer Sequencing)(Hundal et al., 2016; Hundal et al., 2020) following somatic mutation analysis (Figure 1a). PRC2-WT tumours exhibited a slightly higher mutational burden than PRC2-loss tumours (0.82 vs 0.68 mutations/Mb; t-test, p=0.0986, Figure 1b). Stratification by PRC2 complex status revealed comparable total numbers of predicted numbers of pMHC neoantigens between PRC2-WT and PRC2-loss tumours, but with but considerable heterogeneity (median 3 vs 4 per sample; Wilcoxon test, p=0.31; Figure 1c), consistent with similar mutation burdens across subtypes. Surprisingly, 23.9% of PRC2-WT samples lacked any predicted neoantigen derived high-affinity pMHC complexes, whereas 43.5% and 32.6% harboured a range of 1-4 and 5-137 of such neoantigens, respectively. A similar distribution was observed for PRC2-loss (17.8%, 40%, and 42.2%, respectively; Figure 1d, e). Neoantigens from PRC2-WT and PRC2-loss MPNSTs primarily arose from missense variants (97% vs 91.6%), followed by frameshift (3% vs 7.8%) and in-frame deletions (0% vs 0.6%), respectively, displaying a significant difference in variant-type distribution between groups (Fisher’s Exact Test, p=0.0207; Figure 1f).

**Figure 1.**
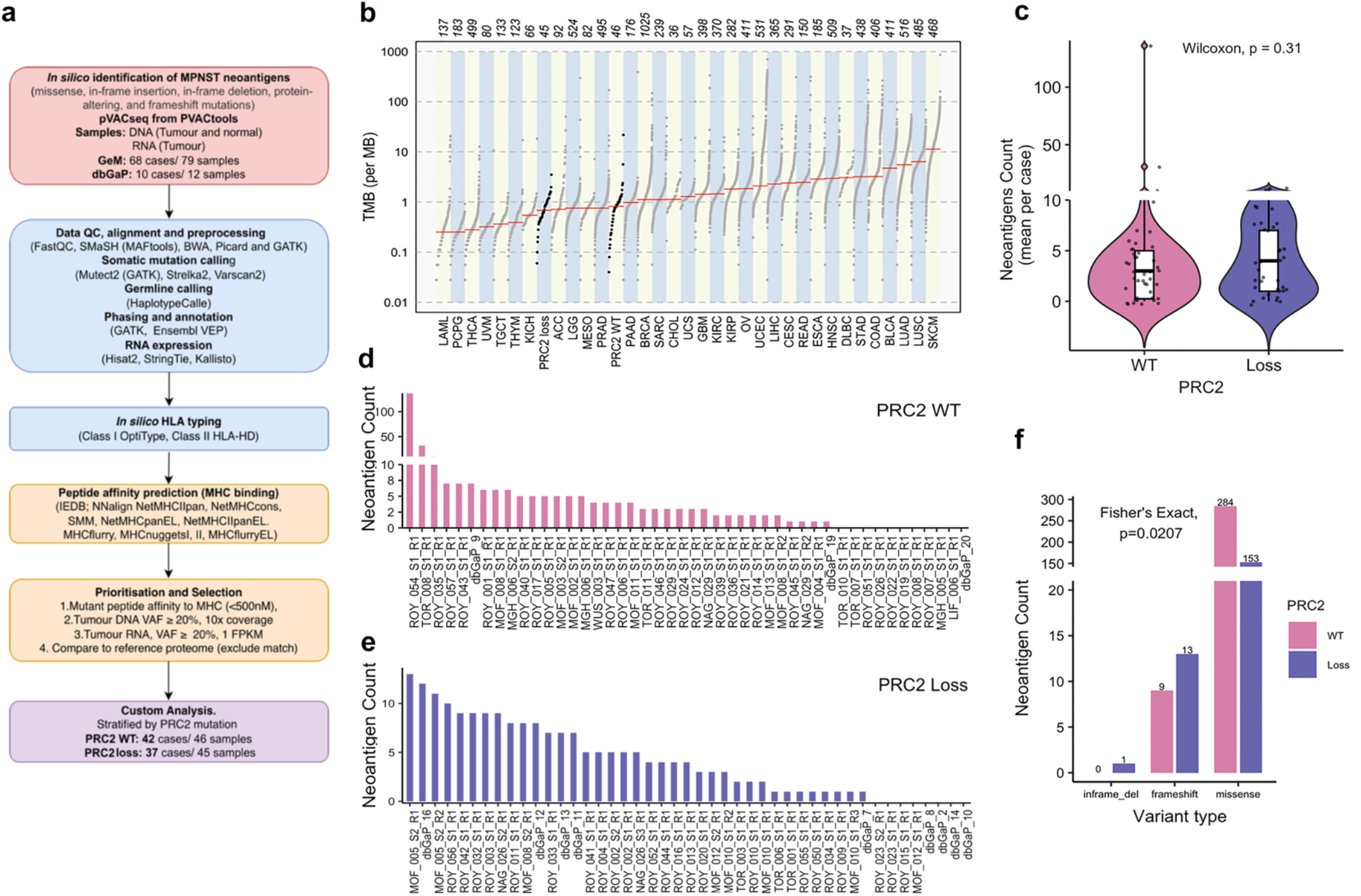
Distribution and characterisation of predicted somatic mutation-derived neoantigens stratified by PRC2 status. **a**. Workflow for identification of somatic neoantigens (missense, in-frame insertion/deletion, protein-altering, and frameshift mutations) using pVACtools with sample inputs (red), pre-processing steps (blue), pVACseq analysis (orange) and PRC2 stratification (purple). All samples passed QC. **b**. Dot plot of MPNST somatic Tumour Mutational Burden (TMB, non-synonymous somatic mutations per megabase) derived from sample WGS and WES (black dots) compared with TCGA reference datasets (grey dots). Numbers displayed indicate the number of samples within the cohort harbouring at least one non-synonymous mutation, red horizontal line median TMB. **c**. Violin plot of the distribution of predicted neoantigens (mean count per case) in PRC2-WT (n=46 samples from 42 cases) and PRC2-loss (n=45 samples from 37 cases), Wilcoxon test. **d**. and **e**. Bar plots displaying the count of the predicted neoantigens across PRC2-WT (d) and PRC2-loss (e) samples. **f**. Bar plots showing the number of predicted neoantigen peptides derived from missense, frameshift, and in-frame insertion/deletion mutations in PRC2-WT and PRC2-loss (Fishers exact test).

Predicted pMHC binding patterns revealed a highly diverse and individualised allelic landscape. In PRC2-WT tumours, 20.5% of neoantigens were predicted to bind MHC class I (60 missense, 1 frameshift) and 79.5% to class II (229 missense, 8 frameshift). The most frequent class I allele was HLA-C*07:02 (four cases, 12 epitopes), followed by HLA-B*07:02 (three cases, 12 epitopes) and HLA-A*02:01 (two cases, 11 epitopes). Among class II alleles, DPA1*01:03 was the most recurrent (eight cases, 22 epitopes), followed by DPB1*02:01 (seven cases, 9 epitopes) and DRB5*01:01, which exhibited the broadest binding spectrum (29 epitopes across three cases; Figure 2a upper panels). In PRC2-loss cases, 16.2% of neoantigens were predicted to bind to class I and 83.8% to class II. HLA-C*12:03 was the most frequent class I allele (four cases, 7 epitopes), followed by HLA-A*68:01 and HLA-C*07:02 (two cases each). Among class II alleles, DPB1*02:01 was most frequent (seven cases, 17 epitopes), followed by DQA1*02:01 (six cases, 14 epitopes), and DPA1*01:03 (six cases, 9 epitopes). Other broad-binding spectrum MHC Class II were DPB1*04:02, and DRB8*01:01, each binding 12 epitopes and observed in four and three cases respectively (Figure 2a, lower panels).

**Figure 2.**
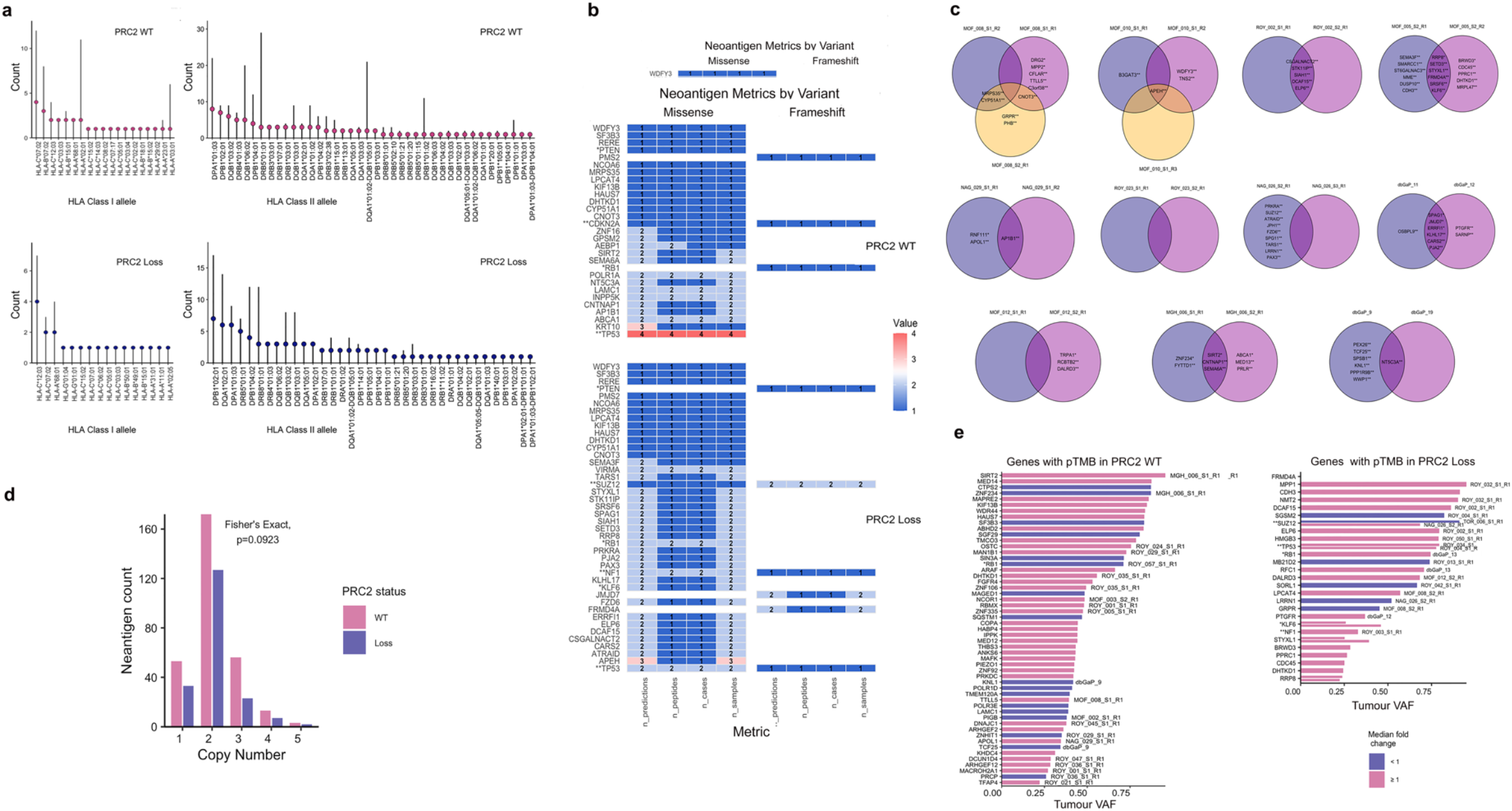
Distribution of predicted neoantigens by HLA allele, gene variant, copy number and persistent TMB, stratified by PRC2 status. **a**. Lollipop plot of predicted peptide counts per individual HLA alleles across the MPNST cohort stratified by PRC2 status. Vertical line represents the number of unique epitopes predicted to bind a given HLA allele. Lollipop head indicates the number of unique MPNST cases carrying that HLA allele. **b**. Heatmaps of specific gene–associated (y-axis) metrics across multiple samples, stratified by variant type (missense or frameshift) and PRC2 status (WT or loss) in 56 genes with more than one prediction. From left to right; n_predictions (the total number of predicted neoantigens for each gene), n_peptides (the total number of unique neoantigen high affinity peptides), n_cases (the number of distinct cases), and n_samples (the total number of samples having the corresponding neoantigen peptide). Numerical values listed highlighted with associated colour gradient). Mutated genes are displayed along the y-axis. ** MPNST driver genes, * tumour suppressor genes. **c**. Venn diagrams summarizing predicted sample heterogeneity for either presence (overlap) or absence of neoantigens (individual samples) detected across multiple samples from the same MPNST case. Overlaps indicate candidates shared between samples; the central intersection denotes neoantigens present in all samples (truncal). Non-overlapping sectors represent sample-restricted candidates. In all the plots pink and purple represent PRC2-WT and PRC2-loss, respectively. **d**. Bar plot of predicted neoantigens by mean case genomic copy number stratified by PRC2 status (Fishers exact test). **e**. Bar plot of genes exhibiting persistent tumour mutation burden (pTMB) against variant allele frequency (VAF) stratified by PRC2 status. Genes are ordered by decreasing VAF. Bars are coloured according to the median fold change (median WT IC_50_ Score / median MT IC_50_ Score): values ≥ 1 are shown in pink, and values < 1 are shown in blue. Split bars represent samples carrying the same mutated gene. Sample identifiers are displayed. Bars without labelling are for samples with highest frequency of neoantigen prediction, ROY_054_S1_R1 in PRC2-WT and for MOF_005 (MOF_005_S2_R1, MOF_005_S2_R2) in PRC2-loss.

### MPNST predicted peptide MHC neoantigens are private

Predicted neoantigens originated from a spectrum of genes, including canonical MPNST drivers, tumour suppressors, and passenger mutations. To refine this landscape, genes with more than one predicted peptide (n=56 genes) were retained and categorised by variant type (missense and frameshift), PRC2 status and the unique epitopes per case and sample. While few genes were recurrently mutated across patients and subgroups (e.g. *TP53*), all the resulting peptides were unique to each case (i.e. private or personalised neoantigens), indicating individualised mutational and antigen-processing profiles. *TP53*-derived neoantigens were detected in four PRC2-WT cases, all from independent missense mutations, with median pMHC IC_50_ values of 11.0-384.5nM (1.3-2.7-fold-change relative to WT; Variant allele frequencies (VAF) =0.50-0.92), each binding prediction to distinct MHC proteins. In PRC2-loss tumours, *TP53* neoantigens occurred in three cases (two missense, one frameshift, none in common with PRC2-WT) with median MT IC_50_ values of 11.3-187.0nM and 2.6-3.6-fold increase over WT peptide. Additional common neoantigen-generating genes included the tumour suppressors, *PTEN, RB1*, and other rarer variants (*DHTKD1, HAUS7, KIF13B, LPCAT4, NCOA6, PMS2, RERE, SF3B3*, WDFY3) (Figure 2b, Supplementary Table 2).

In MPNSTs with PRC2-loss, *NF1*-derived neoantigens were identified in three patients (two missense, one frameshift), with IC_50_ values of 5.7-44.5nM. *SUZ12* mutations (one missense, two frameshift) generated both class I and class II neoantigens with median MT IC_50_ values of 12.1-44.1nM (Figure 2b, Supplementary Table 2) with a VAF ranging from 0.68-0.91. Notably, one *SUZ12* missense mutation derived peptide exhibited a fold change <1 in IC_50_ compared to the wild-type peptide, suggesting it was potentially less antigenic. A *CDKN2A* mutation in a single PRC2-WT case produced two mutant peptides, arising from both missense and frameshift mutations (Figure 2b, Supplementary Table 2).

Substantial intra-patient heterogeneity was also observed in some cases. Multiple samples from the same case shared identical neoepitope sequences predicted to bind the same MHC, indicating consistent antigen presentation within the tumour. However, some samples from the same tumour (sub-clonal regions) also harboured distinct neoepitope sets. For instance, MOF_008 included two PRC2-WT and one PRC2-loss sample, with no shared neoantigens detected across all three samples, though, three mutated gene predicted peptides were common to two of them. Here, *CNOT3* and *MRPS35* produced similar peptides binding the same HLA allele, whereas *CYP51A1* yielded two closely related epitopes -AYVKIKTIWSTMLRL and VKIKTIWSTMLRL-binding DRB5*01:21 and DRB1*15:01, respectively. These differences corresponded to variations in HLA-DH class II typing outputs, where DRB5*01:21 was detected in MOF_008_S2_R1 and MOF_008_S1_R1 but not in MOF_008_S1_R2 (Figure 1i, j). Similar heterogeneity was also observed in MOF_012, where one sample (MOF_012_S1_R1) lacked neoantigens while another (MOF_012_S2_R1) harboured three, and in MGH_006, where two samples shared three common epitopes in addition to private ones (Figure 2c). Together, these data reveal a complex evolution and individualised neoantigen landscape in MPNST, with overlapping candidate mutational driver genes, but no shared specific pMHC predicted epitopes between any of the combined cohort of cases.

### Persistent Tumour Mutational Burden defines a patient-specific higher priority neoantigen repertoire

Persistent tumour mutation burden (pTMB) represents the subset of somatic mutations that are unlikely to be eliminated during tumour evolution, encompassing alterations retained in either single-copy (only-copy) or multi-copy genomic regions. Previous studies have shown that the clinical efficacy of immune checkpoint blockade ICB correlates strongly with these presistent mutations, represented by pTMB, in a lineage-dependent manner (Niknafs et al., 2023). To identify neoantigens arising from persistent mutations, we integrated neoantigen prediction data with copy number profiles (Figure 2d). The overall distribution of copy number states did not differ significantly between PRC2-loss and PRC2-WT tumours (Fisher’s Exact, p=0.923), indicating comparable frequencies of each copy number category across groups.

In the PRC2-WT cohort, 53 neoantigens were associated with genes in only (single)-copy regions across 17 samples, and 73 neoantigens were linked to multi-copy genes across 16 samples (Figure 2e, left panel). These neoantigens appeared strong immunogenic candidates, as they were expressed at the RNA level, of clonal origin (all with DNA VAF ≥ 0.2), and frequently exhibited high variant allele fractions. Specifically, 35.8% of only-copy and 24.7% of multi-copy neoantigens had VAFs >0.5. Although observed across several cases, more than half of these originated from ROY_054_S1_R1, the sample with the highest overall neoantigen load. Most of these peptides also showed mutant-to-wild-type median IC_50_ (fold changes >1, suggesting stronger predicted MHC binding for the mutant epitopes.

Predicted higher quality pTMB neoantigens from persistent tumour mutations were also identified in PRC2-loss, as 33 neoantigens were associated with genes in only-copy across 17 samples, 18 (17 missense and 1 frameshift) of them with VAF >0.5, and 13 had IC_50_ fold changes >1 (Figure 2e, right panel). Multi-copy genes contributed 32 neoantigens across 20 samples, 7 (6 missense and 1 frameshift) that had VAF >0.5 and 5 have IC_50_ fold changes >1. Taken together, these findings demonstrate that both PRC2-WT and PRC2-loss MPNSTs harboured a subset of persistent, clonal, and transcriptionally expressed neoantigens-arising from both single-copy and multi-copy genomic regions-with mutant-specific MHC binding characteristics, supporting the biological relevance of pTMB and prioritisation of these as high-confidence candidates for individualised immunotherapeutic strategies despite the overall low neoantigen burden.

### Patient-Specific Fusion-derived Neoantigen Landscape and HLA Diversity

For this analysis, the PRC2-WT group was composed of 45 cases (76 samples) and the PRC2-loss was from 38 cases (58 samples) (Figure 3a). Gene fusions represent a valuable source of tumour-specific antigens due to their capacity to generate novel open reading frames and entirely non-self-peptide sequences at the fusion breakpoints. We evaluated the repertoire of fusion-derived neoantigens in MPNST stratified by PRC2 status using tumour RNA sequencing. The overall number of predicted fusion-derived neoantigens appeared comparable between PRC2-WT and PRC2-loss tumours (median 5 vs 6 per sample; Wilcoxon, p=0.69; Figure 3a) yet, there was a marked heterogeneity within both groups.

**Figure 3.**
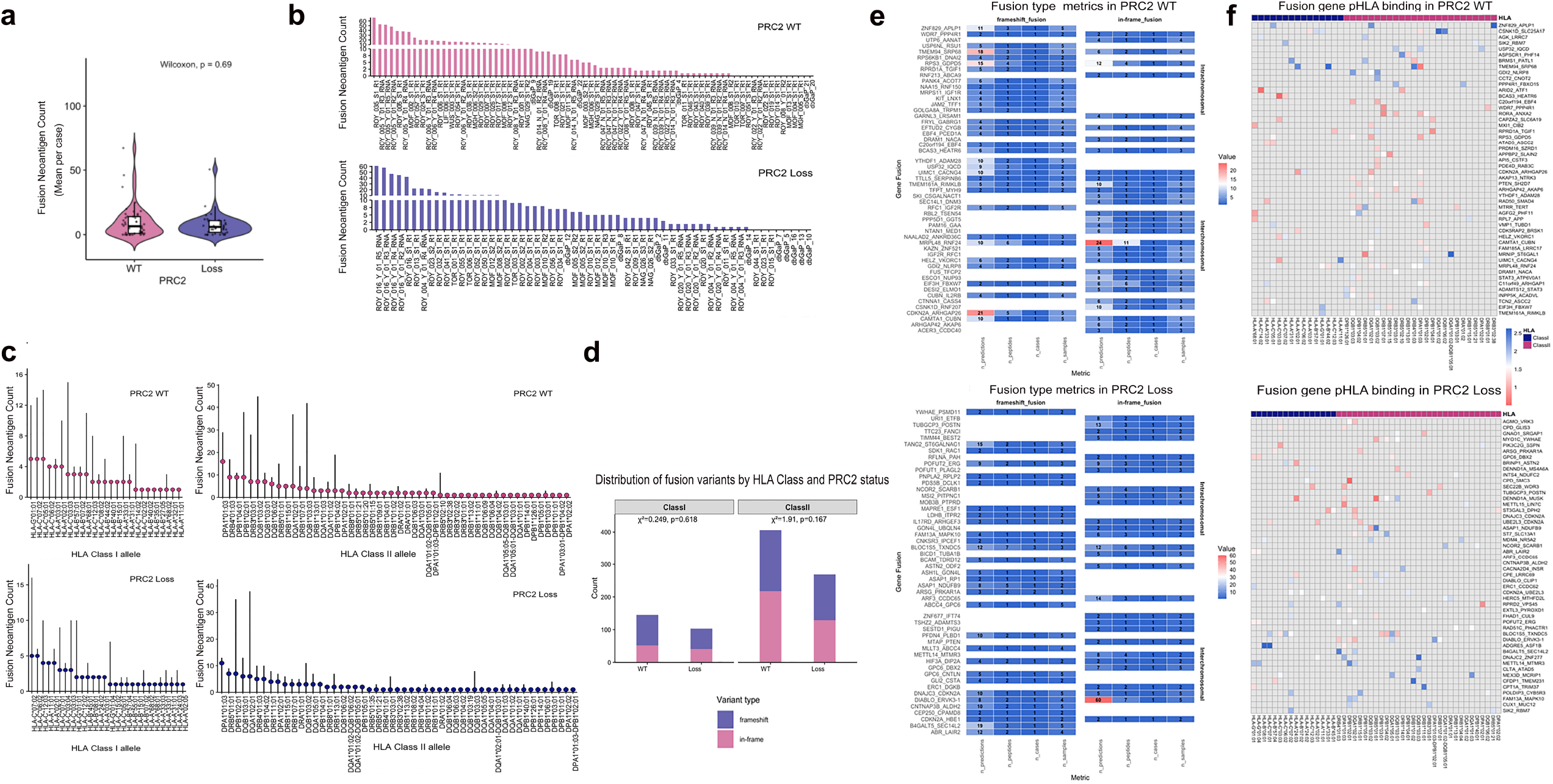
Distribution and characterisation of fusion-derived neoantigens stratified by PRC2 status. **a**. Violin plots of predicted gene fusion derived neoantigens per MPNST case using pVACfuse stratified by PRC2 status. PRC2-WT (76 samples from 45 cases) and PRC2-loss tumours in blue (58 samples from 38 cases), Wilcoxon test. **b**. Bar plot displaying the count of the predicted unique fusion derived neoantigens across all samples as in (a). **c**. Lollipop plots of predicted fusion gene neoantigen counts with respect to individual HLA alleles across the cohort cases. Each vertical line represents the number of unique neoantigens predicted to bind a given HLA allele. The head of the lollipop indicates the number of unique cases carrying that allele. **d**. Stacked bar plots showing the number fusion-derived neoantigen peptides originating from frameshift (blue) and in-frame (pink) stratified by PRC2 status and HLA class, Pearson’s Chi-squared test. **e**. Heatmaps of specific fusion–associated (y-axis) metrics across multiple samples, stratified by variant type (frameshift or inframe), fusion type (intra-chromosomal or inter-chromosomal) and PRC2 status (PRC2-WT= 52 fusion genes, or PRC2-loss = 50 fusion genes). From left to right; n_predictions (the total number of predicted fusion neoantigens for each gene), n_peptides (the total number of unique fusion neoantigen high affinity peptides), n_cases (the number of distinct cases), and n_samples (the total number of samples having the corresponding fusion neoantigen peptide). Numerical values listed highlighted with associated colour gradient. Fusion genes are displayed along the y-axis. **f**. Heatmaps of the binding affinity median score (log_10_ IC_50_) for predicted peptide-HLA interactions among the top 50 HLA alleles and top 50 fusion genes stratified by PRC2 status. Rows represent fusion genes and columns correspond to HLA alleles grouped by class (I and II). All HLA alleles shown exhibit strong binding affinities (IC_50_ ≤ 500 nM). Colour; red indicates stronger predicted binding (lower IC_50_ values) compared to blue.

In PRC2-WT tumours, 873 predicted neoantigens corresponded to 548 unique epitopes from 331 fusion genes. Most samples carried few neoantigens: 18.4% had no predicted neoantigen derived high affinity pMHC complexes, 50.0% between 1-10, 22.4% between 11-30, and 9.2% more than 30. Seven samples (from three cases) accounted for 45.6% of all predicted neoantigens (Figure 3b, upper panels). The PRC2-loss group showed a similar distribution, with 366 unique epitopes from 196 fusion genes; 13.7% of samples lacked predicted neoantigen derived high affinity pMHC complexes, 56.9% had up to ten, 20.7% is from 11-30 and 8.6% exceeded 30 -all from a single case (ROY_016), which contributed 43.8% of the total neoantigen load (Figure 3b, lower panels). Within individual cases, unique fusion neoantigen counts were typically consistent across samples, being uniformly high or low. For example, ROY_005 (5 samples, 38-53 neoantigens), ROY_016 (5 samples, 42-61 neoantigens), and ROY_047 (5 samples, 2-3 neoantigens). Overall, these observational data suggested comparable fusion-derived neoantigen burdens between subgroups but with inter-tumoural heterogeneity.

A total of 52 HLA alleles were shared between the PRC2-WT and PRC2-loss groups, and all were predicted to bind fusion-derived neoantigens, indicating a broadly overlapping HLA repertoire between the cohorts (Figure 3c). In both groups, the class II allele DPA1*01:03 was the most frequent (PRC2-WT: 16 cases, 29 distinct epitopes, PRC2-loss: 11 cases, 13 epitopes). The class II allele DQA1*02:01 exhibited the greatest peptide-MHC binding range (PRC2-WT: seven cases, 45 epitopes, PRC2-loss: six cases,38 epitopes). DQB1*03:02 was the MHC to bind 42 unique epitopes in 7 cases in the PRC2-WT compared to 11 unique epitopes in 3 PRC2-loss cases (Figure 3c, supplementary Table 3). Similarities were also seen for class I alleles: HLA-C*07:02 was the commonest class I allele (five cases in each group) and the MHC recognised 14 and 16 unique epitopes in PRC2-WT and PRC2-loss, respectively. HLA-C*06:02 was observed in four PRC2-WT and PRC2-loss cases, the MHC recognising four to six unique epitopes. HLA-C*05:01 was more frequent in WT (five cases) and recognised 14 different epitopes compared to 1 case with 2 different epitopes in PRC2-loss group (Figure 3c, supplementary Table 3). Certain alleles were group-specific, including HLA-B*57:01 (three WT cases) and HLA-C*07:01, HLA-C*03:04, and DQA1*05:01 (three mutant cases each, recognising four - nine epitopes). Eighteen alleles presented a single unique epitope across distinct cases (Figure 3f, g, supplementary Table 3).

Fusion-derived neoantigens originated from both frameshift and in-frame fusion variants in PRC2-WT and PRC2-loss tumours. These variants showed no significant difference in distribution between the two groups (frameshift: WT=281, loss=202; in-frame: WT=270, loss=170; χ^2^=0.843, p=0.359) (Figure 3d). Both HLA class I and II molecules contributed similarly to predicted presentation. In PRC2-WT tumours, class I neoantigens comprised 10.1% (frameshift) and 5.6% (in-frame), while class II represented 20.4% and 23.6%, respectively. In PRC2-loss tumours, the corresponding values were 4.4% and 6.7% for class I, and 14.0% and 15.2% for class II. Differences in class distribution were not significant (class I: χ^2^=0.249, p=0.618; class II: χ^2^=1.91, p=0.167; Figure 3d).

### Unique Fusion-genes per case with distinct predicted peptide repertoires

Despite the above findings, many of the fusion events are in fact non-validated and likely to be artefactual and non-functional. Many were excluded from additional downstream analysis to avoid overinterpretation of their as yet unclear significance, and to maintain stringent criteria for neoantigen candidate selection (Supplementary Table 4). For most fusion genes, either inter- or intra-chromosomal in origin, these were unique to individual cases producing distinct predicted peptide repertoires. A small number of fusion genes occurred in both frameshift and in-frame forms - for example, POFUT2_ERG, BLOC1S5_TXNDC5 and DIABLO_ERVK3-1 in PRC2-loss tumours, and UIMC1_CACNG4 and TTLL5_SERPINB6 in PRC2-WT tumours-yielding different peptide sequences that bound distinct HLA alleles (Figure 3e). As part of our initial filtering, we excluded non-tumour fusion events, including recurrent artefacts and fusions commonly detected in normal tissues (Methods, supplementary Table 4). One intra-chromosomal fusion, OR51S1_TP53I11, was detected in 17 cases across 24 samples (11 PRC2-WT, 6 PRC2-loss) and initially yielded 23 predicted neoantigens corresponding to three unique epitopes; one epitope was shared among 21 samples (Supplementary Figure 2a, b). OR51S1_TP53I11 has previously been reported only once in colorectal cancer (Jeon et al., 2022) and could be a novel fusion gene in MPNST. We visually inspected the putative breakpoints in the Integrative Genomics Viewer (left breakpoint chr11:4,848,669; right breakpoint chr11:44,933,050). No clear evidence of a breakpoint was observed and the supporting reads were suggestive of alignment artefacts (Supplementary Figure 3). We further assessed this event using FusionInspector, a component of the Trinity Cancer Transcriptome Analysis Toolkit (CTAT) that re-evaluates STAR-Fusion predictions.

FusionInspector failed to validate OR51S1_TP53I11 in any sample, instead listing it in the failed_reads_during_span_analysis file and flagging it with a “low_per_id” status, indicative of low sequence identity and insufficient read support. Consequently, non-validated fusion events were excluded. Three samples, ROY_022_01_R1, ROY_008_Y_01_R2, and ROY_022_Y_01_R2_RNA lost their neoantigens with this filter and 521 unique fusion genes across the cohort were identified. Only eight were shared between PRC2-WT and PRC2-loss tumours, reflecting limited overlap in the fusion landscape.

Among shared fusions, BLOC1S5_TXNDC5 generated 13 epitopes (seven from frameshift and six from in-frame events) across three mutant cases (IC_50_=5.15 to 136.9nM, binding class I and class II HLA alleles). It is considered a read-through transcript regulated by nonsense-mediated decay. ARSG_PRKAR1A (PRC2-loss) genes are located next each other and produced two epitopes from frameshift events (IC_50_=4.7-10.45nM, binding class I and class II alleles) (Figure 4). Three fusion genes; GOLGA8A_TRPM1, SIK2_RBM7 and SLC36A4_SIK3, were shared between groups within the same case (MOF_008), which included two PRC2-WT and one PRC2-loss samples. MOF_008 harboured ten fusion genes in total; four unique to the PRC2-loss sample and three unique to the PRC2-WT, illustrating intra-patient fusion heterogeneity (Supplementary Table 3).

**Figure 4.**
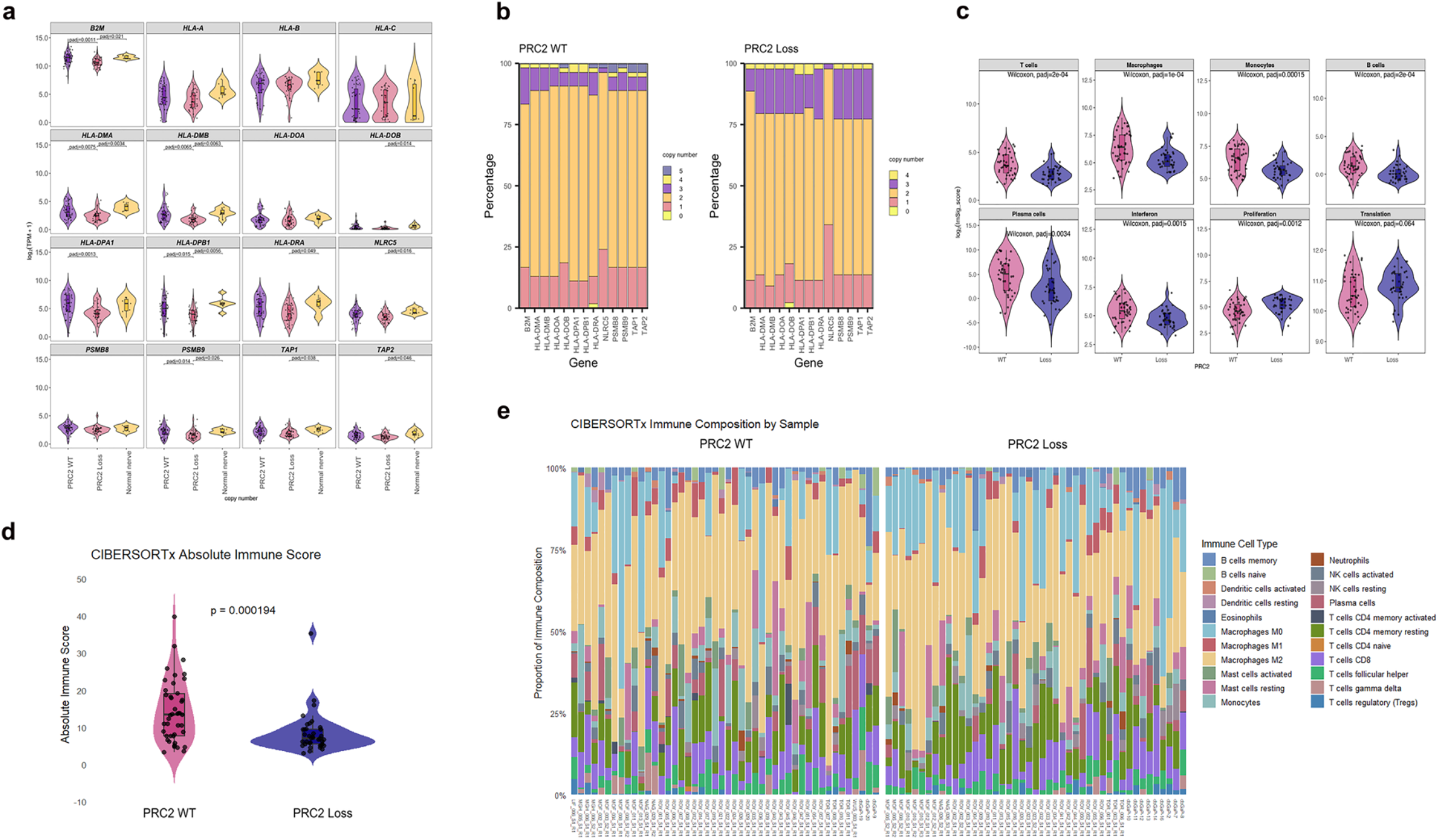
Transcriptome of MPNST antigen presentation and immune microenvironment stratified by PRC2 status. **a**. Violin plots antigen-processing gene expression (mean expression per case, log_2_(TPM+1)) stratified by PRC2 status; PRC2-WT (n=46 samples from 42 cases), PRC2-loss (n=45 samples from 37 cases), and normal nerve (n=7). Statistical analysis using Kruskal-Wallis test, followed by Dunn Test for multiple comparisons with Holm adjustment. Significant comparisons are displayed. Within each violin, box plots indicate the median and interquartile range (IQR), whiskers represent 1.5 times the IQR, and jittered points show individual sample measurements. **b**. Stacked bar plots showing the percentage of the average genomic copy number per case for antigen processing and presentation pathway genes stratified by PRC2 status; (PRC2-WT, n=54 patients from 61 samples) (PRC2-loss, n=44 patients from 56 samples). Statistical significance evaluated using Fisher’s Exact Test and with adjusted p-value using FDR. **c**. Violin plot of the ImSig-score, reflecting cell abundance score of immune cells and biological process, stratified by PRC2 status. Wilcoxon rank-sum followed with adjusted p-values using FDR. **d**. Absolute immune infiltration scores in PRC2 WT and PRC2 Loss samples, estimated by CIBERSORTx using bulk-RNA sequencing data. Violin plots with overlaid boxplots are shown along with individual data points. Replicates from the same sample, but different regions were averaged before computing statistical significance using a two-tailed, unpaired, Wilcoxon rank sum test. e. Relative composition of immune cell subsets inferred from bulk RNA-sequencing data using CIBERSORTx with the in-built LM22 immune-cell gene signature matrix. Stacked bar plots display the proportional composition of 22 immune cell subsets, shown as percentages of total immune cell infiltration, for each sample, grouped by PRC2 status (PRC2 WT and PRC2 Loss), see also Heatmap in Supplementary Figure 8.

Fusion genes frequently generated multiple predicted neoantigens, producing several unique peptides per fusion in both groups. Predicted epitopes were also broadly consistent across different samples from the same case (Figure 3f). For example, in PRC2-WT the fusion CDKN2A_ARHGAP26 (five samples from one case) yielded 21 predictions and five unique peptides; these peptides were predicted to bind DPA1*01:03, DQA1*02:01 and HLA-A*02:01, and HLA-C*06:02 (IC_50_=8.521-61.46nM). RPS3_GDPD5 (three samples from one case) produced 27 predictions (frameshift and in-frame) and eight unique peptides (IC_50_=7.15-40.57nM) for DPA1*01:03, DQA1*02:01, DQB1*03:03 and HLA-C*05:01 (Figure 4a, c, Supplementary Table 3). Similarly, in the PRC2-loss group DIABLO_ERVK3-1 (five samples from ROY_016 case) yielded 68 predicted fusion epitopes and nine distinct peptides predicted to bind six class II alleles (IC_50_=5.29 and 159.1nM) (Figure 3f, Supplementary Table 3). This fusion gene contributed to 10.1% of all the predicted neoantigens in the PRC2-loss group. Taken together, these results demonstrate that while fusion-derived neoantigen burden and the HLA allelic landscape are broadly similar between MPNST molecular subtypes, the neoantigen distribution is highly uneven across individual cases and that the pathogenic fusion-gene repertoires exhibit no overlap between them.

### Neoantigen burden and prognostic relevance

Having characterised neoantigen load across MPNST subtypes and both cohorts, we next assessed whether there was any prognostic relevance of neoantigen burden on patient outcome. Neoantigens predicted by pVACseq and pVACfuse were aggregated at the sample level and averaged across multiple samples per patient (Supplementary Figure 4). The overall neoantigen burden was comparable between PRC2-WT and PRC2-loss tumours, with final median values of 10 and 11 neoantigens per sample, respectively. Stratification of cases into low (≤5), medium (6-15), and high (≥16) neoantigen load groups revealed no significant differences in overall survival (OS) in either PRC2-WT tumours (log-rank p=0.37) or PRC2-loss tumours (log-rank p=0.26). When the entire cohort was analysed jointly, neoantigen burden was not significantly associated with OS (log-rank p=0.22). However, this analysis remains underpowered, and the limited cohort size restricts definitive conclusions regarding any prognostic impact of neoantigen load (Supplementary Figure 4b).

### PRC2 loss is associated with suppression of antigen-processing machinery and reduced immune infiltration in MPNST

Having identified both somatic and fusion-derived neoantigens and characterised their distribution across PRC2-WT and PRC2-loss tumours, we next independently evaluated whether the cellular machinery required for antigen-processing and presentation was preserved in these tumours. As effective neoantigen presentation depends on coordinated peptide processing and MHC class I and II expression for CD8^+^ and CD4^+^ T-cell recognition, we analysed the transcriptional landscape of key antigen-presentation genes to assess the impact of PRC2 status. Specifically, we evaluated RNA expression of MHC class I components class I components (*HLA-A, HLA-B, HLA-C*, and *B2M*), peptide-processing regulators (*PSMB8, PSMB9, TAP1, TAP2, NLRC5*), and MHC class II molecules (*HLA-DMA, HLA-DMB, HLA-DOA, HLA-DOB, HLA-DPA1, HLA-DPB1, HLA-DRA*) in MPNST tumours and normal nerve tissue (Figure 4a). Overall, significant group effects were observed for multiple antigen-presentation genes, particularly MHC class II components (Kruskal-Wallis, p<0.05; Supplementary Table 5a). Pairwise comparisons demonstrated significantly higher expression of *B2M* (1.8-fold), *HLA-DMB* (2.0-fold), *HLA-DMA* (1.6-fold), *PSMB9* (1.9-fold), *HLA-DPA1* (3.8-fold), and *HLA-DPB1* (2.0-fold) in PRC2-WT tumours compared with PRC2-loss, with similar reduced expression relative to normal nerve tissue (Figure 4a). *NLRC5, TAP1, TAP2, HLA-DRA*, and *HLA-DOB* also showed reduced expression in PRC2-loss tumours compared with normal nerve. The coordinated downregulation of multiple MHC class II genes suggests impaired CD4^+^ T-cell priming and helper function, further constraining effective anti-tumour immunity in PRC2-loss tumours. In contrast, MHC class I genes (*HLA-A, HLA-B, HLA-C*) did not show significant expression differences between PRC2-WT and PRC2-loss tumours. Moreover, no significant expression differences were observed between normal nerve and PRC2-WT tumours for any antigen-presentation gene (Dunn’s test, padj<0.05; Figure 4a).

We next assessed copy number alterations in antigen-presentation genes. Copy number states for *HLA-A, HLA-B*, and *HLA-C* could not be evaluated because this highly polymorphic region exhibited insufficient mappable coverage, resulting in exclusion of these bins during CNV calling. Among the remaining genes, no significant differences in the distribution of mean copy number states (gain, neutral, loss) were observed between PRC2-WT and PRC2-loss tumours (Fisher’s exact test, FDR<0.05; Supplementary Table 5b). Neutral copy number predominated in both groups (66.6-77.2% in PRC2-WT; 61.3-77.2% in PRC2-loss), while one-copy loss ranged from 13.0-24.1% in PRC2-WT and 9.1-34.1% in PRC2-loss tumours. NLRC5 exhibited the highest frequency of one-copy loss, occurring in 34.1% of PRC2-loss cases (Figure 4b). In addition, a single missense mutation with moderate predicted impact in *NLRC5* was identified in one tumour (ROY_023_01), with no recurrent damaging mutations detected in other antigen-presentation genes. Taken together, although classical MHC class I genes were preserved, suppression of *NLRC5, TAP1/2, B2M* and immunoproteasome components in PRC2-loss tumours would have been sufficient to impair peptide processing and MHC class I loading, thereby compromising effective antigen presentation.

Effective anti-tumour immunity also requires immune cell infiltration and functional cytotoxic T-cell responses within the tumour microenvironment. We next characterised immune composition and effector gene expression across MPNST subtypes first using the ImSig deconvolution framework (Nirmal et al., 2018). Of 569 ImSig marker genes, 549 were detected in our dataset, with 463 retained following dataset-specific feature selection based on co-expression (rho≥0.6). Most immune and functional modules demonstrated strong gene coverage and co-expression, supporting their reliability for abundance scoring. Following feature selection, strong correlations were observed for macrophage (median rho=0.75), T-cell (rho=0.65), monocyte (rho=0.6), plasma cell (rho=0.75), and interferon-signalling modules (rho=0.57) (Figure 4c). In contrast, neutrophil and NK-cell signatures showed weaker co-expression (median rho<0.4), suggesting limited or heterogeneous representation of these populations (Supplementary Table 5c). Abundance scores were calculated as the median expression of retained marker genes, and downstream analyses were restricted to modules with moderate to strong co-expression.

When comparing PRC2-WT and PRC2-loss tumours, significant differences were observed across most immune modules. PRC2-WT tumours exhibited coordinated enrichment of T cells (1.75-fold), macrophages (2.5-fold), monocytes (1.9-fold), B cells (2.1-fold), and plasma cells (11.7-fold), together with increased interferon-signalling activity (1.7-fold), relative to PRC2-loss tumours (Wilcoxon test, FDR < 0.05). In contrast, PRC2-loss tumours were characterised by significantly higher proliferation scores (1.5-fold; FDR < 0.05), with no significant difference in translation-associated modules (Figure 4c, Supplementary Table 5b). These findings confirm that PRC2 status is a major determinant of the immune microenvironment in MPNST.

To complement the ImSig analysis, we also examined expression of key immune activation and effector genes. Consistent with T-cell module enrichment, *CD4, LCK*, and *IL2RA* were significantly upregulated in PRC2-WT tumours, with median increases of 2.1-fold, 2.3-fold, and 7.8-fold, respectively, compared with PRC2-loss tumours (Dunn’s test, padj<0.05). Expression of the chemokines *CCL2* and *CCL5* was also significantly lower in PRC2-loss tumours, showing 2.2-fold and 3.5-fold higher median expression in PRC2-WT tumours, respectively (Dunn’s test, padj<0.05; Supplementary Figure 5). Although *CD8A, CD8B*, and *FOXP3* were higher in PRC2-WT tumours, these differences did not reach statistical significance. In contrast, cytotoxic effector genes *PRF1, GZMB*, and *GZMA* were significantly upregulated in PRC2-WT tumours (≈2.3-fold). Markers of monocyte and macrophage lineages (*CD14, CD33, ITGAM/CD11b*) were also significantly higher in PRC2-WT tumours (1.9-2.3-fold), supporting a broader reduction in myeloid infiltration in PRC2-loss tumours (Supplementary Figure 5, Supplementary Table 5e).

**Figure 5.**
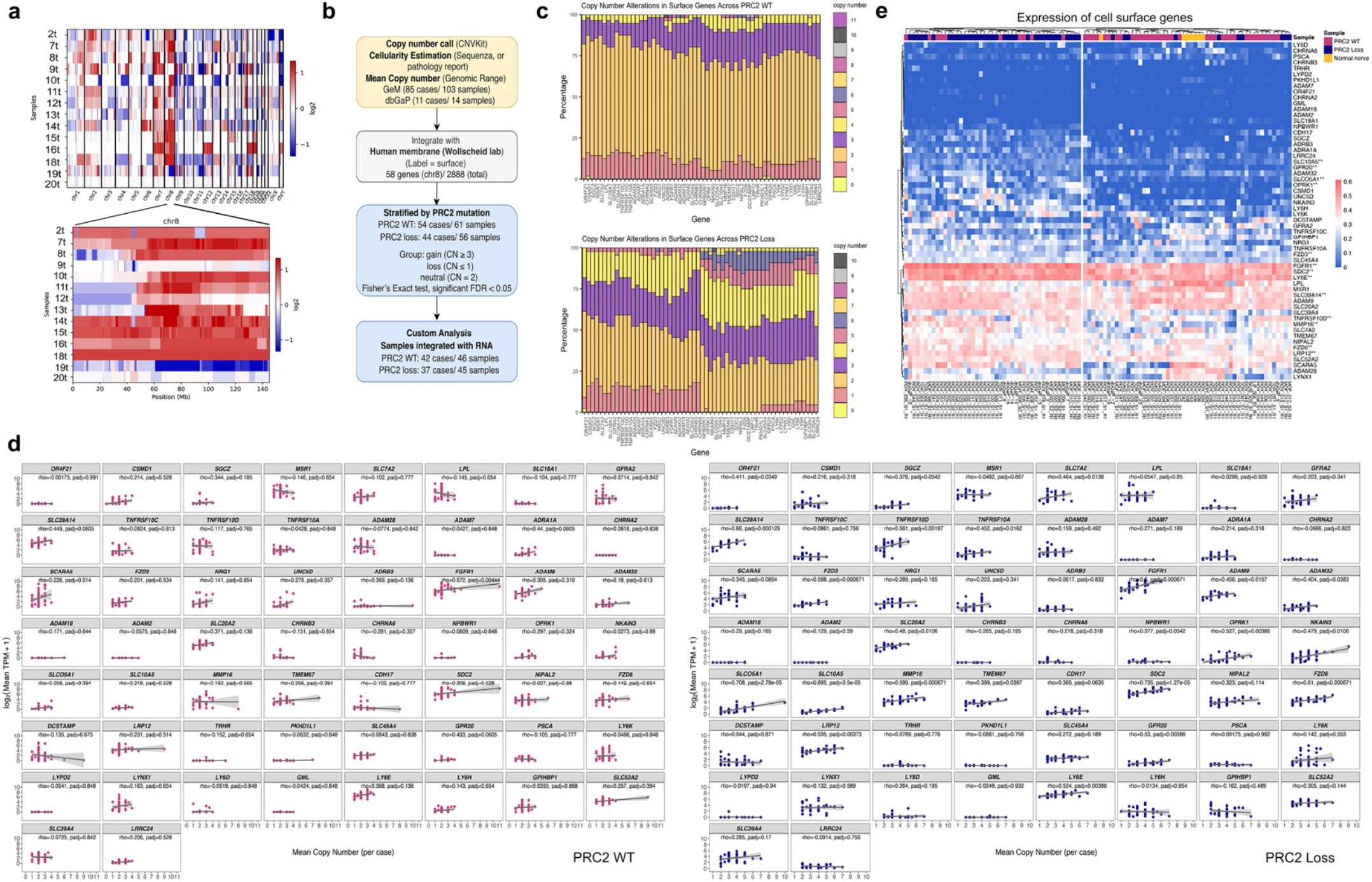
Genomic copy number dependent overexpression of candidate MPNST cell surface antigens stratified by PRC2 status. **a**. Heatmap of log2 ratio of chromosomal copy number for all chromosomes in the MPNST dbGaP cohort (n=14) (top) and for chromosome 8 (below). Red, white and blue indicate copy number gain, neutral and loss. Note: some samples arise from the same case, 9t with 19t, 11t with 12t, 7t and 18t. Heatmaps generated using CNVkit. **b**. Workflow to identify copy number dependent cell surface proteins as potential MPNST immune targets. **c**. Stacked bar plots of the frequency of the mean copy number (per case) for chromosome 8 cell surface genes stratified by PRC2 status (PRC2-WT n=54 cases, PRC2-loss group n=44). Genes (x-axis) are arranged by chromosome 8 position. **d**. Dot plots of correlations between copy number and mRNA expression for each candidate MPNST cell surface gene across chromosome 8, log_2_(TPM+1) stratified by PRC2 status. For cases with multiple tumour samples, expression values were averaged to derive a single mean value per case. Spearman correlation coefficients (rho) p-value adjusted using FDR. Grey shaded plot region is the 95% confidence interval around the regression line. **e**. Clustered heatmap of gene expression the expression, log_10_(TPM+1), of chromosome 8 surface genes in MPNST cohorts and normal nerve expression (n=7, yellow). Red and blue, high and low gene relative gene expression. For tumour cases with multiple samples, expression values were averaged per case. Rows represent genes and columns represent samples. Genes marked with double asterisks (**) show a moderate to strong correlation between copy number and RNA expression (rho >0.5, FDR <0.05) in PRC2-loss tumours. Both genes and samples were hierarchically clustered using Euclidean distance and complete-linkage.

Next, we assessed inhibitory immune checkpoint signalling. Expression of *CD274* (PD-L1) was significantly reduced in PRC2-loss tumours with 2.6-fold higher median expression in PRC2-WT; Dunn’s test, padj<0.05), while *PDCD1* (PD-1) showed more modest differences, with a 3.5-fold higher median expression in PRC2-WT tumours (padj=0.045). KLRG1, an inhibitory receptor associated with terminally differentiated cytotoxic lymphocytes, was also upregulated in PRC2-WT tumours (3.1-fold; Dunn’s test, FDR <0.05, Supplementary Figure 5, Supplementary Table 5e). No significant differences were detected for either *CTLA4, LAG3*, or *CD160*.

We next complemented this analysis by using immune cell deconvolution with CIBERSORTx and both the dbGAP and GEM bulk-RNA-Seq samples (Figure 4d, e). Here, we utilised the LM22 leukocyte gene signature matrix and inferred the abundance of 22 immune cell subsets across individual samples stratified by PRC2 status (Supplementary Figure 8). These data show the general higher infiltration of M2 Macrophages, monocytes CD8 and CD4 memory T-cells in the PRC2 WT compared with the PRC2-loss samples, consistent with a higher overall absolute immune score (Figure 4d, p=0.00019, Wilcoxon). Moreover, this is further represented by the relative proportions of immune composition across samples (Figure 4e).

Collectively, these results indicate that immune suppression in PRC2-loss MPNST is not driven by enhanced checkpoint-mediated inhibition, but rather reflects a global reduction in immune cell infiltration and cytotoxic effector activity, consistent with an immune-desert phenotype compared to PRC2-WT.

### Copy number gain-associated expression changes in chromosome-8 surface genes in PRC2-loss MPNST

Recurrent chromosomal gains represent a prominent source of tumour-specific transcriptional dysregulation in MPNST. In the dbGaP cohort, chromosome 8 copy number gain was observed in 78.6% of cases (11 cases from 14 samples), with six samples exhibiting whole-chromosome gain and five showing partial gains predominantly involving the long arm of chromosome 8 (chr8q) (Figure 5a). This pattern recapitulates recurrent chr8q amplification previously reported in MPNST and closely associated with H3K27me3 loss and PRC2 deficiency (Cortes-Ciriano et al., 2023). Consistent with this association, all cases harbouring chromosome 8 gain in the dbGaP cohort carried PRC2 mutations (Lee et al., 2014). Together, these findings establish chromosome 8 -particularly chr8q-as a recurrently amplified genomic region in PRC2-loss MPNST, providing a rationale for subsequent analyses of chromosome 8 encoded surface genes with potential relevance for TAA discovery that may be inform immunotherapy strategies.

To identify candidate TAAs, we integrated the GeM and dbGaP cohorts and stratified patients by PRC2 status (PRC2-WT vs PRC2-loss) (Figure 5b). Copy number states were assigned at the patient level (gain, cn≥3; neutral, CN=2; loss, cn≤1) using mean values across samples from the same patient. The distribution of copy number states differed significantly between the two groups for 57 chromosome 8 ‘surface’ genes, with the exception of *OR4F21* (Fisher’s exact test; FDR < 0.05; Figure 5b; Supplementary Table 6b). In PRC2-WT, the gain was observed in 11.1% to 25.9% for surface genes in chromosome 8 short arm (chr8p) and from 22.2% to 35.2% in chr8q. The percentage of the gain was higher in the PRC2-loss, with 38.6% to 56.8% in chr8p and in chr8q the gain was seen between 59.1% to 75% of the samples (Figure 5c, Supplementary Figure. 5a, Supplementary Table 5a). Sample-level copy number profiles were visualised as heatmaps, revealing largely consistent copy number states across samples from the same patient, with only limited intra-tumoural heterogeneity (Supplementary Figure. 5c).

We next correlated mRNA expression with copy number across cases stratified by PRC2 status. While in PRC2-WT tumours (42 cases corresponding to 46 samples) only FGFR1 reached a moderate effect (Spearman rho=0.572, FDR=0.004; Figure5d left panel, Supplementary Table 5c), PRC2-loss tumours showed 13 genes with moderate to strong effects (rho≥0.5, FDR <0.05; Figure 5d, right panel, Supplementary Table 5). These included four genes on chr8p (*SLC39A14, TNFRSF10D, FZD3*, and *FGFR1*) and nine on chr8q (*OPRK1, SLCO5A1, SLC10A5, MMP16, SDC2, FZD6, LRP12, GPR20*, and *LY6E*). Together, these findings highlight a recurrent set of chromosomes 8-primarily chr8q-encoded surface genes whose expression is associated with genomic amplification which is characteristic of MPNST tumours with H3K27me3 loss as a consequence of PRC2 loss of function.

We then compared mRNA expression in tumour samples with normal nerve tissue from GeM cohort (n=7). Normal nerve demonstrated lower expression and clustered primarily with PRC2-WT samples and the majority of the PRC2-loss clustered together in a heat map (Figure. 5e). In PRC2-WT tumours, comparison of gene expression across copy number states revealed modest differences for five genes (Kruskal-Wallis test, p<0.05). Because these analyses were restricted to a limited set of pre-specified genes, p-values were not adjusted for multiple comparisons, consistent with established statistical guidance (Rothman, 1990). However, following pairwise multiple-comparison only *ADRA1A* and *FGFR1* showed significant difference between tumours with copy number gain and neutral with median fold of 0.96 and 2.1 higher in gain, respectively (Dunn’s test, padj<0.05, Supplementary Figure7a). In contrast, PRC2-loss tumours exhibited a higher frequency of copy number gain, with many genes lacking representation in the loss category (29 genes; no loss cases for 26 genes and only a single loss case for three genes). Consequently, in addition to three group analyses, direct comparisons between copy number gain and neutral tumours were performed. This revealed multiple genes with significant upregulation in the gain group. Notably, *SLCO5A1, SLC10A5, GPR20*, and *SDC2* demonstrated median expression increases ranging from 1.5 to 2.17-fold in gained tumours relative to neutral (Wilcoxon test, FDR<0.05). *LY6E, MMP16, FZD6, OPRK1*, and *SLC39A4* also exhibited substantial gain-associated increases (≥1.8-fold), although these effects were more modest in statistical significance (Wilcoxon test, p<0.05; Supplementary Figure 6). Moreover, genes represented across all three copy number states displayed a similar dosage-dependent pattern. In particular, *FGFR1* and *SLC39A14* showed increases of approximately 2.5-fold and 1.5-fold, respectively, in the gained tumours compared to neutral (Kruskal-Wallis test followed by Dunn’s test, padj<0.05; Supplementary Figure7b, lower panel). An additional eight genes showed more modest gain-associated expression differences, with fold changes ranging from 1.8 to 3.1 (Kruskal-Wallis test, p<0.05; Supplementary Table 6). The expression of copy number gain between the PRC2-loss and PRC2-WT tumours was not statistically significant for all chromosome 8 surface genes apart from *SLC10A5, GPR20*, and *NKAIN3*, in which the expression was more for the PRC2-loss gain was statistically significant (Wilcoxon test, FDR<0.05, Supplementary Figure7c). Together, these analyses identify a recurrent set of chromosome-8 specially chr8q encoded, amplification-associated surface genes that are consistently overexpressed in PRC2-loss MPNST, nominating candidate tumour-associated antigens for further functional and preclinical evaluation.

## Discussion

MPNSTs remain highly aggressive sarcomas with limited effective systemic therapies, underscoring the urgent need for new treatment strategies, including immunotherapy. To our knowledge, this is the first study that integrated genomic and transcriptomic data of MPNSTs to characterise computationally predicted tumour antigenicity. The results demonstrate that personalised neoantigens (private) appear to predominate across both PRC2-WT and PRC2-loss tumours. Moreover, PRC2-loss tumours exhibit substantial reduction in the expression of components of the antigen processing and presentation machinery together with globally reduced immune infiltration, consistent with an immune-cold tumour microenvironment. In contrast, these tumours show recurrent overexpression of several cell-surface TAAs associated with copy number gain of chromosome 8, potentially highlighting an alternative and complementary immunotherapeutic opportunity. Together, these tumour NGS derived predictive findings support an antigen-centred framework in which personalised neoantigen repertoires and overexpressed surface TAAs may be leveraged in a PRC2-dependent manner to broaden immunotherapeutic strategies for MPNST.

Given the rarity of MPNST and the limited availability of high-quality tumour material, this study relied on restricted-access public sequencing datasets generated by the GeM Consortium (Cortes-Ciriano et al., 2023) and dbGaP (Lee et al., 2014). There was neither immune-proteomic data available nor validated TCR sequencing data. To identify tumour-specific antigens, we applied state-of-the-art neoantigen computational prediction using updated versions of the pVACseq and pVACfuse pipelines from pVACtools (Hundal et al., 2016; Hundal et al., 2019; Hundal et al., 2020). These tools integrate multiple biologically relevant features inot trained machine learning algorithms that try and characaterise the neoantigen landscape and to prioritise high-confidence neoantigens. These tools address mutant-specific MHC binding relative to the corresponding wild-type peptide, consideration of nearby germline variants that may influence binding, reference proteome similarity filtering is used to exclude self-like sequences, and confirmation of tumour RNA expression. We further applied stringent filtering for fusion-derived neoantigens, excluding candidates detected in normal tissues or lacking independent fusion validation. In parallel, gene copy number driven expression analysis enabled the identification of candidate TAAs, providing a complementary antigen class for therapeutic consideration.

Consistent with previous NGS reports, MPNSTs fall into two main groups, those that either retain or lose PRC2 repressor complex function. Despite this classifier, an overall low tumour mutational burden, including persistent tumour mutational burden, was observed across both groups compared to melanoma and other solid tumours, and both somatic mutation-derived and fusion-derived neoantigen loads were at a modest level, with a median total of approximately 10 private neoantigens per tumour. Although a wide range of neoantigen counts were observed, the upper extreme was largely driven by a single outlier case, underscoring the overall low numbers of neoantigens in most tumours. These features are consistent with emerging clinical and preclinical evidence suggesting that checkpoint inhibition as a single agent appears unlikely to provide consistent long-term benefit in MPNSTs, at least based on TMB assessment alone (Lingo et al., 2025), likely reflecting limited tumour immunogenicity and anti-tumour neoantigen priming rather than dominant checkpoint-mediated immune suppression. For MPNSTs, results of ICB therapies have not resulted in universal long-term clinical responses despite the detection of immune inflammatory infiltration in PRC2-WT compared to PRC2-Loss associated MPNSTs. For instance, in a cohort of 53 MPNST patients, only 7/53 (13%) had at least PD-L1 labelling (5%), and 57% had TILs (CD8, 5%) labelling using immunohistochemistry (Shurell et al., 2016). Microarray expression data for six MPNST tumours showed PDL-1 expression to be downregulated compared to plexiform neurofibroma, the precursor of MPNST, and CTLA-4 appeared upregulated (Haworth et al., 2017). This heterogeneity appears to be mainly associated with the PRC2 complex, as loss of PRC2 function promoted immune evasion and an immune-desert tumour microenvironment (Yan et al., 2022). Moreover, MPNST patient samples showing an immune inflammatory infiltrate in tumours are frequently those associated with an intact PRC2 complex (Cortes-Ciriano et al., 2023). Direct injection of a modified vaccinia virus Ankara (MVA) enhances tumour immunity, a strategy that can alter the immune-desert TME, and appeared to sensitise the PRC2-Loss MPNSTs to ICB therapy (Wu et al., 2021). ICB therapy alone, however, was only effective in a few cases of MPNSTs that showed PDL-1 expression (Davis et al., 2019; Larson et al., 2022; Ozdemir et al., 2019). These data suggest that immune therapy in the appropriate context in MPNST may be only partially effective.

Despite the low neoantigen burden, accumulating evidence indicates that alternative immune-based strategies, particularly cancer vaccines, can be effective in tumours with limited mutational loads in order to generate a polyclonal T-cell response. Personalised neoantigen vaccination has demonstrated strong and sustained antigen-specific T-cell responses in pancreatic ductal adenocarcinoma, a typical low-TMB and immune-cold tumour, in the context of a prospective clinical trial conducted in the adjuvant setting (Rojas et al., 2023). Also, vaccination targeting the single, clonal IDH1-R132H neoantigen in IDH-mutant glioma elicited sustained mutation-specific T-cell responses in a first-in-human phase I trial, providing clinical proof-of-principle that effective vaccination can be achieved despite low levels neoantigen presentation (Platten et al., 2021). Across tumour types, vaccine efficacy has been shown to depend mainly on neoantigen quality rather than quantity, including clonality, transcriptional expression, persistence, and favourable mutant-specific MHC binding properties (Niknafs et al., 2023; Rojas et al., 2023). Moreover, clinical studies consistently indicate that optimal responses are achieved in settings of minimal residual disease and/or in combination with strategies that enhance T-cell priming and trafficking (Lang et al., 2022; Ott et al., 2018), providing important design principles for future vaccine-based immunotherapy trials in MPNST.

A defining feature of PRC2-loss MPNSTs, as reported by several other groups, includes profound suppression of immune cell infiltration, including both lymphoid and myeloid populations. Genomic and transcriptomic studies have shown that PRC2 inactivation is associated with downregulation of antigen processing and presentation machinery, attenuated interferon signalling, and reduced immune infiltration, resulting in an immune-desert tumour microenvironment (Cortes-Ciriano et al., 2023; Yan et al., 2022; Zhang et al., 2024). Here, transcriptomic subtyping further supports this distinction, identifying PRC2-deficient tumours as immune-low subgroups characterised by reduced macrophage and T-cell signatures and poorer clinical outcomes (Holand et al., 2023). In this context, immunotherapeutic strategies that rely on endogenous antigen presentation may be intrinsically constrained in MPNST.

Recurrent copy number gain affecting chromosome 8 in MPNSTs represent a prominent genomic feature of PRC2-loss MPNST and provide a biologically grounded source of TAA overexpression. We identified a recurrent, dosage-dependent upregulation of multiple chromosome-8-encoded surface genes, including *FGFR1, SDC2, SLCO5A1, GPR20, LY6E*, and *MMP16*, with minimal intra-tumoural heterogeneity across samples from the same patient. PRC2-loss MPNST, where antigen presentation and immune cell recruitment are compromised, TAA-directed CAR-T cell therapies offer a potential rational alternative by enabling direct, MHC-independent tumour targeting (Brudno et al., 2024; Zugasti et al., 2025). In this circumstance, adoptive cellular therapy (ACT) strategies include based on the ex-vivo stimulation and expansion of T cells expressing selective TCRs or CARs, either by genetic modification (monoclonal) or by expansion of tumour infiltrating lymphocytes (polyclonal) before infusion back into patients (Qian and Liu, 2025; Zhao et al., 2022). While our findings clearly do not establish therapeutic efficacy, they are predictions only, but provide impetus to evaluate set of amplification-driven surface antigens that warrant functional validation and preclinical evaluation as potential targets for TAA-directed cellular therapies.

Several limitations should be acknowledged with predictive computational analysis. Firstly, cryptic neoantigens resulting from peptides generated from short-RNAs would not necessarily be identified in our current approach. These may be identified in the future by using stringent long read allele specific RNA sequencing and immune-peptidomic approaches. Neoantigen prioritisation here was only based on *in silico* predictions of MHC binding affinity and mutant-specific binding advantage, which still remain imperfect surrogates of antigen presentation and immunogenicity. Functional validation of immunogenicity using ELISPOT assays were not feasible in this retrospective analysis due to the lack of matched patient blood PBMC samples in the NGS patient cohorts established by others. Moreover, this study focused on neoantigens derived from somatic mutations and gene fusions and did not assess other potential antigenic sources, such as aberrant splicing. Despite these limitations, our findings provide initial proof-of-principle that MPNSTs can harbour tumour-specific neoantigens that could be subsequently identified and validated, supporting the development of personalised private antigen-centred immunotherapeutic strategies for this disease such as mRNA vaccine in an adjuvant settings or CAR T cell therapy.

## Supporting information

Supplementary Tables Description

## Author contributions

MS performed all data analysis, wrote the paper and generated figures, JC managed the computational cluster, supported software installation and package testing. QQ supported fusion gene analysis, MR performed CIBERSORTX and generated associated figures. BH secured funding, conceived the research plan, wrote and edited the manuscript.

## Acknowledgements

We thank Susanna Kiwala and Malachi Griffith with help with PVACTools (Washington University School of Medicine), the numerous members of Genomics of MPNST Consortium, Jenny and Luke Grenfell-Shaw for funding and strategic support, Oxford data infrastructure (COSMIC and BMRC) and services in the Sir William Dunn School of Pathology.

