## Supplementary Figures for "Tumour neoantigen repertoire computational predictions in malignant peripheral nerve sheath tumours define potential targets for immunotherapy"

**Supplementary Figure 1.**
**Study cohorts and Case Genetic fingerprinting.**

**Supplementary Figure 2.**
**Fusion gene landscape and predicted peptide-HLA binding affinities in PRC2-WT and PRC2-loss.**

**Supplementary Figure 3.**
**Visualisation of the *OR51S1_TP53I11* fusion in the Integrative Genomics Viewer (IGV).**

**Supplementary Figure 4.**
**MPNST overall survival and neoantigen burden.**

**Supplementary Figure 5.**
**Downregulation of immune marker gene expression and immune checkpoint stratified by PRC2**

**Supplementary Figure 6.**
**Copy number alterations of chromosome 8 stratified by PRC2 status in MPNST.**

**Supplementary Figure 7.**
**Copy number alterations and expression patterns of chromosome 8 cell surface genes stratified by PRC2 status in MPNST.**

**Supplementary Figure 8.**
**CIBERSORTx immune cell subset heatmap with LM22 immune-cell signature matrix.**

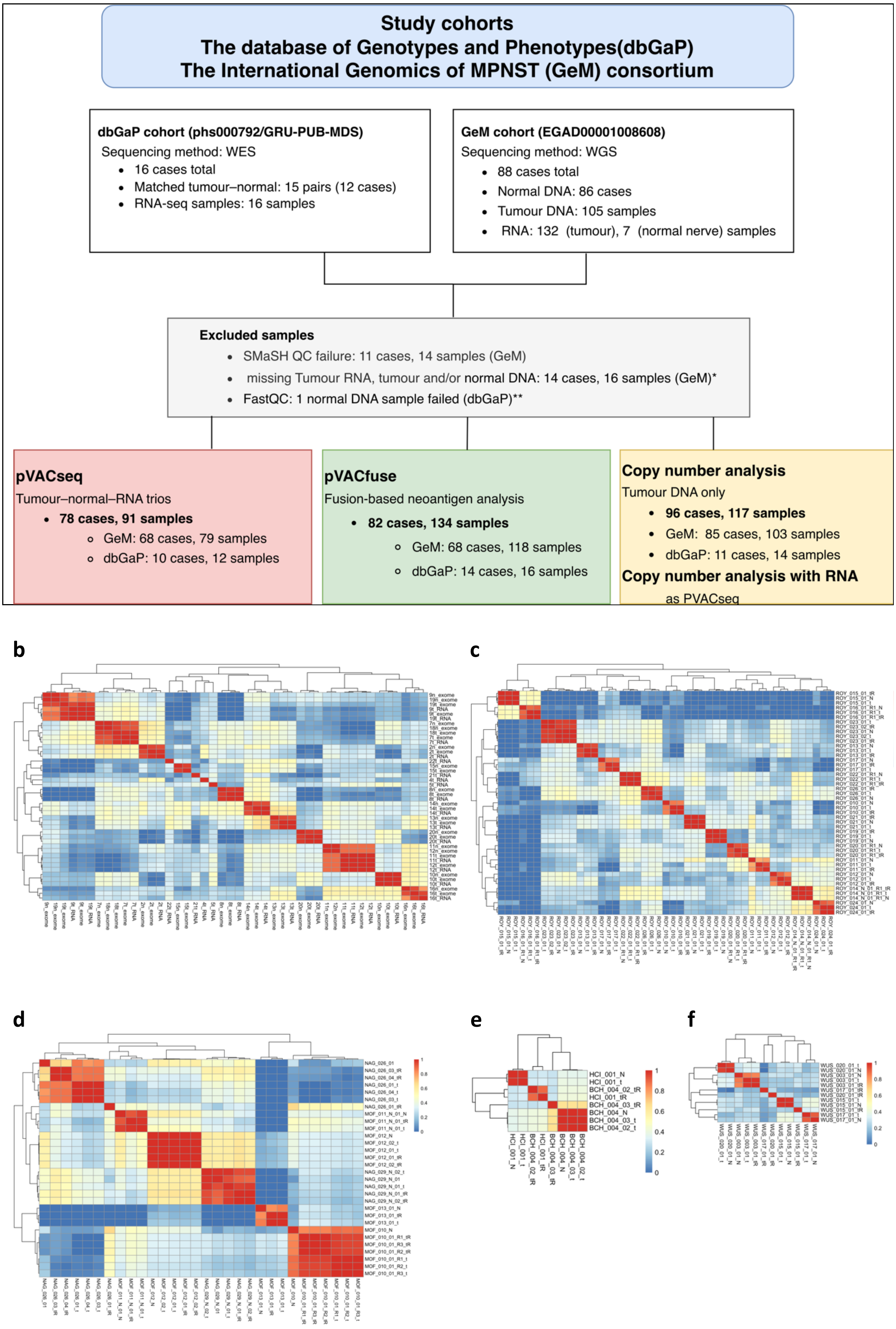
Study cohorts
The database of Genotypes and Phenotypes(dbGaP)
The International Genomics of MPNST (GeM) consortium
dbGaP cohort (phs000792/GRU-PUB-MDS)
Sequencing method: WES
16 cases total
Matched tumour–normal: 15 pairs (12 cases)
RNA-seq samples: 16 samples
GeM cohort (EGAD00001008608)
Sequencing method: WGS
88 cases total
Normal DNA: 86 cases
Tumour DNA: 105 samples
RNA: 132 (tumour), 7 (normal nerve) samples
Excluded samples
SMaSH QC failure: 11 cases, 14 samples (GeM)
missing Tumour RNA, tumour and/or normal DNA: 14 cases, 16 samples (GeM)*
FastQC: 1 normal DNA sample failed (dbGaP)**
pVACseq
Tumour–normal–RNA trios
78 cases, 91 samples
GeM: 68 cases, 79 samples
dbGaP: 10 cases, 12 samples
pVACfuse
Fusion-based neoantigen analysis
82 cases, 134 samples
GeM: 68 cases, 118 samples
dbGaP: 14 cases, 16 samples
Copy number analysis
Tumour DNA only
96 cases, 117 samples
GeM: 85 cases, 103 samples
dbGaP: 11 cases, 14 samples
Copy number analysis with RNA
as PVACseq
b
c
d
e
f

**Supplementary Figure 1.**
**Study cohorts and Case Genetic fingerprinting.**

**a**. Flow chart of samples, cohorts, exclusion criteria and analysis. Cases represent unique patients and may comprise one or more tumour samples, samples refer to individual tumour specimens used for sequencing analyses. **b-e.** Representative heatmaps showing pairwise correlation matrices of SNP VAFs calculated across samples. Each cell represents the Pearson correlation coefficient of VAFs between two BAM files, with colour intensity reflecting the degree of correlation (blue =low, red =high). High correlation values along the diagonal indicate matching genetic fingerprints, consistent with correctly paired samples from the same individual. Heatmaps illustrating correlated samples from the same case in the dbGaP (**b**) and GeM (**c)** cohorts. In the dbGaP dataset, samples sharing the same numerical identifiers correspond to matching exome and RNA pairs (tumour RNA). **d**. Example from case NAG_026, where six of seven samples show strong correlation, whereas NAG_026_01_tR does not match, suggesting a possible mismatch. **e-f.** Examples of unmatched samples in the GeM cohort, including WUS_020, WUS_017, and WUS_015, samples HCI_001 and BCH_004. Samples from the same GeM case share the same three-character prefix followed by a three-digit identifier (e.g., “NAG_026”), denoting case-level grouping.

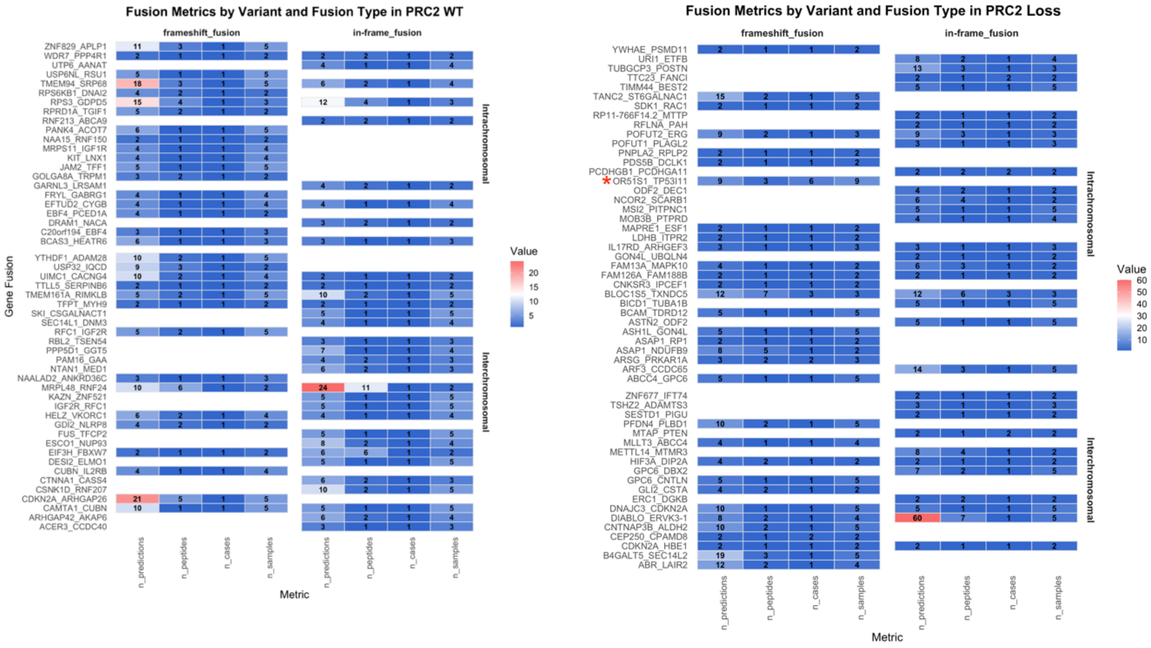


**Supplementary Figure 2.**

**Fusion gene landscape and predicted peptide-HLA binding affinities in PRC2-WT and PRC2-loss.**

Heatmaps illustrating the distribution of fusion gene associated metrics across multiple samples, stratified by variant type (frameshift or in-frame), fusion type (intra-chromosomal or inter-chromosomal) and PRC2-WT (left, n=59 fusion genes) and PRC2-loss (right, n=55 fusion genes), before excluding failed fusion genes that failed validation with FusionInspector. All shown metrics are summarised by fusion gene, variant type, and fusion type, and include:n_predictions (the total number of predicted neoantigens within each group), n_peptides (the total number of unique epitopes), n_cases (the number of distinct cases), and n_samples (the total number of samples having the corresponding epitope). Numerical values within each cell denote the counts for these metrics. The colour gradient represents increasing values, with blue indicating lower and red indicating higher counts. Fusion genes are displayed along the y-axis. *OR51S1_TP53I11* fusion gene is marked with red star.

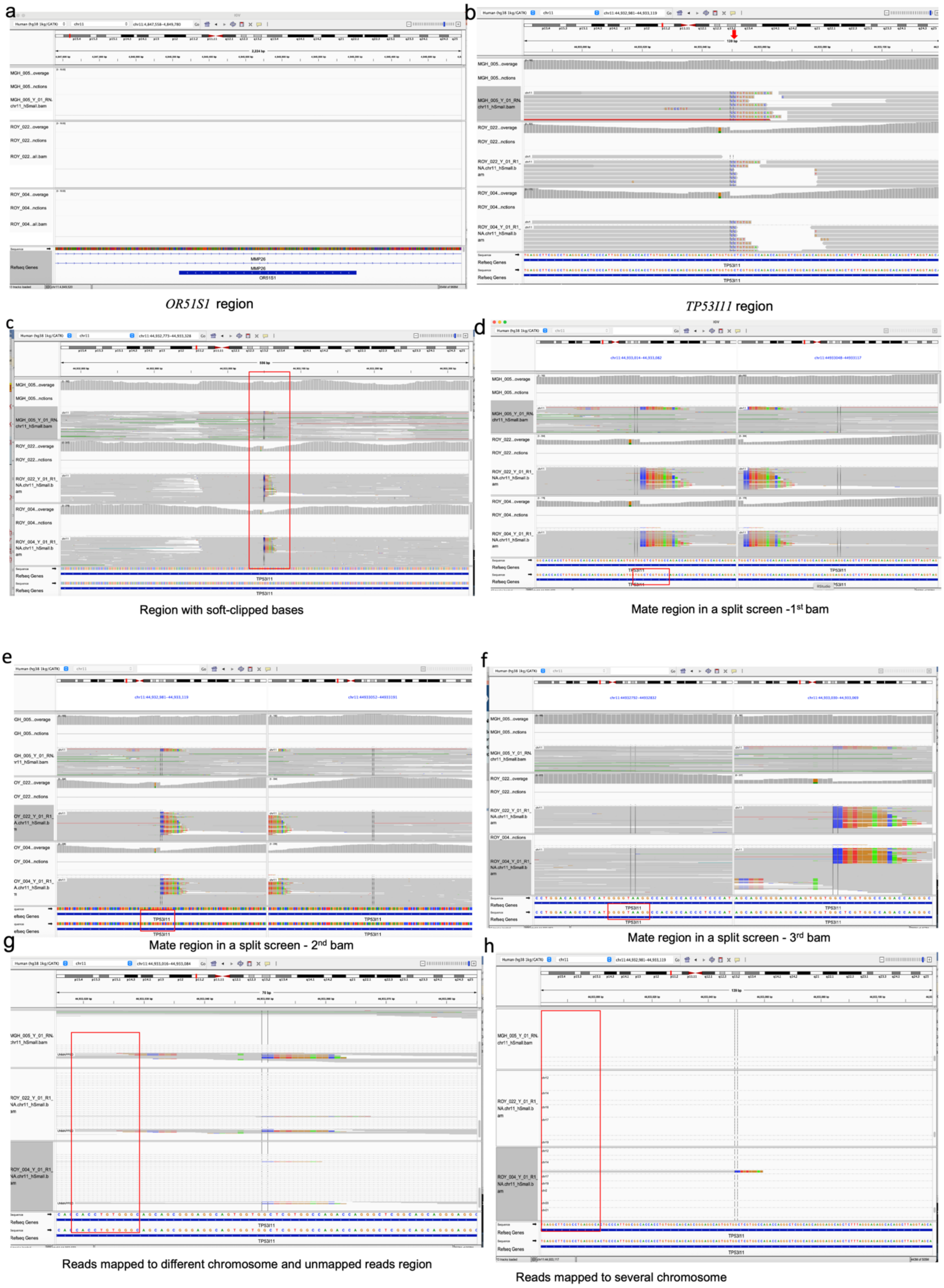


**Supplementary Figure 3.**

**Visualisation of the *OR51S1_TP53I11* fusion in the Integrative Genomics Viewer (IGV).**

**a.** IGV view of the *OR51S1* genomic region showing the putative left breakpoint at chr11:4,848,669. No aligned reads are observed at this site, and zooming out reveals the full

*OR51S1* gene context. **b.** The corresponding right breakpoint region at chr11:44,933,050 within *TP53I1*1 (arrow). Reads are displayed with soft-clipped bases enabled and sorted by base, with alignments grouped by mate chromosome and coloured by insert size and pair orientation. **c.** Zoomed-out view of the *TP53I11* locus showing multiple reads with soft-clipped bases in the second and third BAM tracks, indicative of potential fusion-supporting reads (red box). **d-f**. Split-screen views showing representative mate reads from the soft-clipped regions in the first, second, and third BAM files, each mapping to *TP53I11*. **g-h**. Regions showing reads that align to different chromosomes or remain unmapped (red box). The squished view (*g*) and expanded view (**h**) reveal that several reads in the second and third BAM files map to multiple chromosomes. The first BAM file corresponds to a sample lacking the *OR51S1_TP53I11* fusion, whereas the second and third BAM files represent PRC2-WT and PRC2-loss samples, respectively, both containing the predicted *OR51S1_TP53I11* fusion.

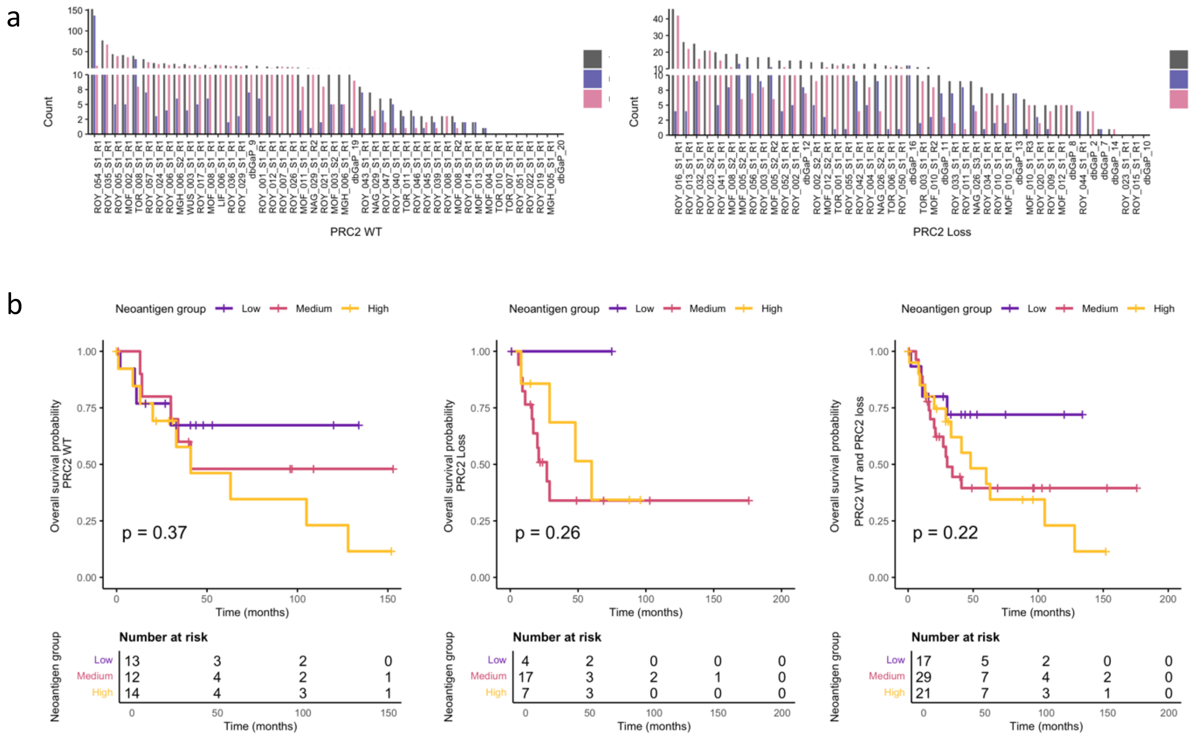


**Supplementary Figure 4.**

**MPNST overall survival and neoantigen burden.**

**a**. Summary bar plot showing the number of predicted neoantigens per case, stratified by PRC2 mutation status. For each case, bars represent the total number of neoantigens (grey), neoantigens derived from somatic point mutations identified by the pVACseq pipeline (blue), and neoantigens derived from fusion genes identified by the pVACfuse pipeline (pink). Only cases that were successfully processed through both pipelines are shown. Cases are displayed separately for PRC2-WT (left) and PRC2-loss (right) groups. **b.** Patients were stratified into low (≤5), medium (6-15), and high (≥16) neoantigen burden groups. Kaplan-Meier curves were generated using the survfit function, and survival differences were assessed by log-rank test. Corresponding numbers at risk are shown beneath the plot. The survival data was only available for the GeM cohort. Only samples that have both pVACseq and pVACfuse data were utilised. Right panel PRC2-WT (n=37), middle panel PRC2-loss (n=27), right panel, combined (n=64).

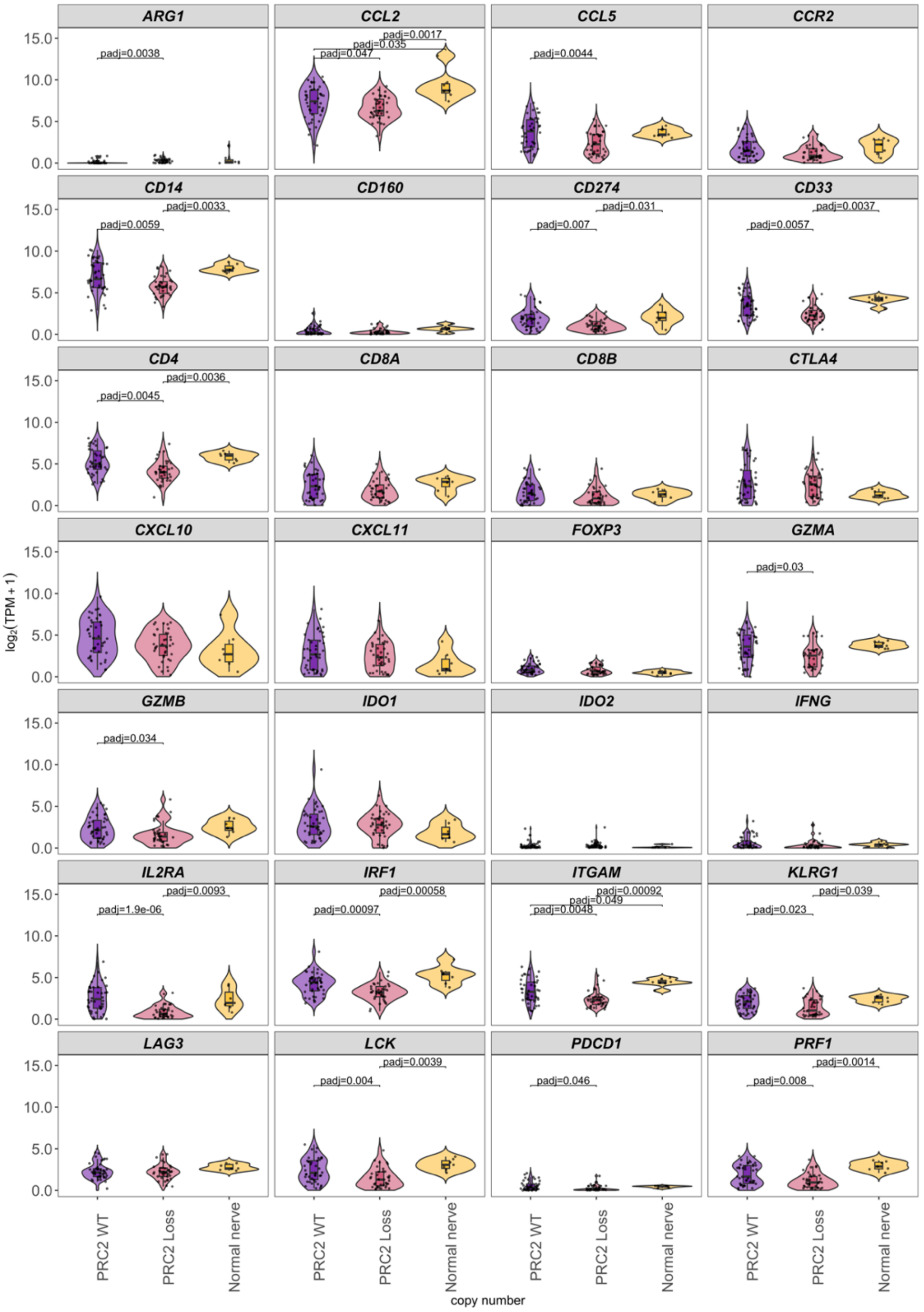


**Supplementary Figure 5.**

**Downregulation of immune marker gene expression and immune checkpoint stratified by PRC2**

Violin plots of antigen-processing gene expression in MPNST cohorts; PRC2-WT (n=46 samples from 42 cases, purple), PRC2-loss (n=45 samples from 37 cases, pink), and normal nerve (n=7, yellow). Statistical significance (mean expression per case) using Kruskal-Wallis test, followed by Dunn test for multiple comparisons and Holm adjustment. Significant comparisons are displayed. Within each violin, box plots indicate the median and interquartile range (IQR), whiskers represent 1.5 times the IQR, and jittered points show individual sample measurements. Y-axis represent the $\log_2$(TPM+1). Immune recruitment and interferon signalling (*CCL2, CCL5, CXCL10, CXCL11, CCR2, IRF1, IFNG*), T-cell abundance and activation (*LCK, CD4, CD8A, CD8B*), cytotoxic effector function (*PRF1, GZMA, GZMB*), regulatory T-cell markers (FOXP3, IL2RA), immune checkpoint and inhibitory receptors (*PDCD1, CD274, CTLA4,*

*LAG3, CD160, KLRG1*), metabolic immune suppression (*IDO1, IDO2, ARG1*), and myeloid lineage markers (*ITGAM, CD33, CD14*).

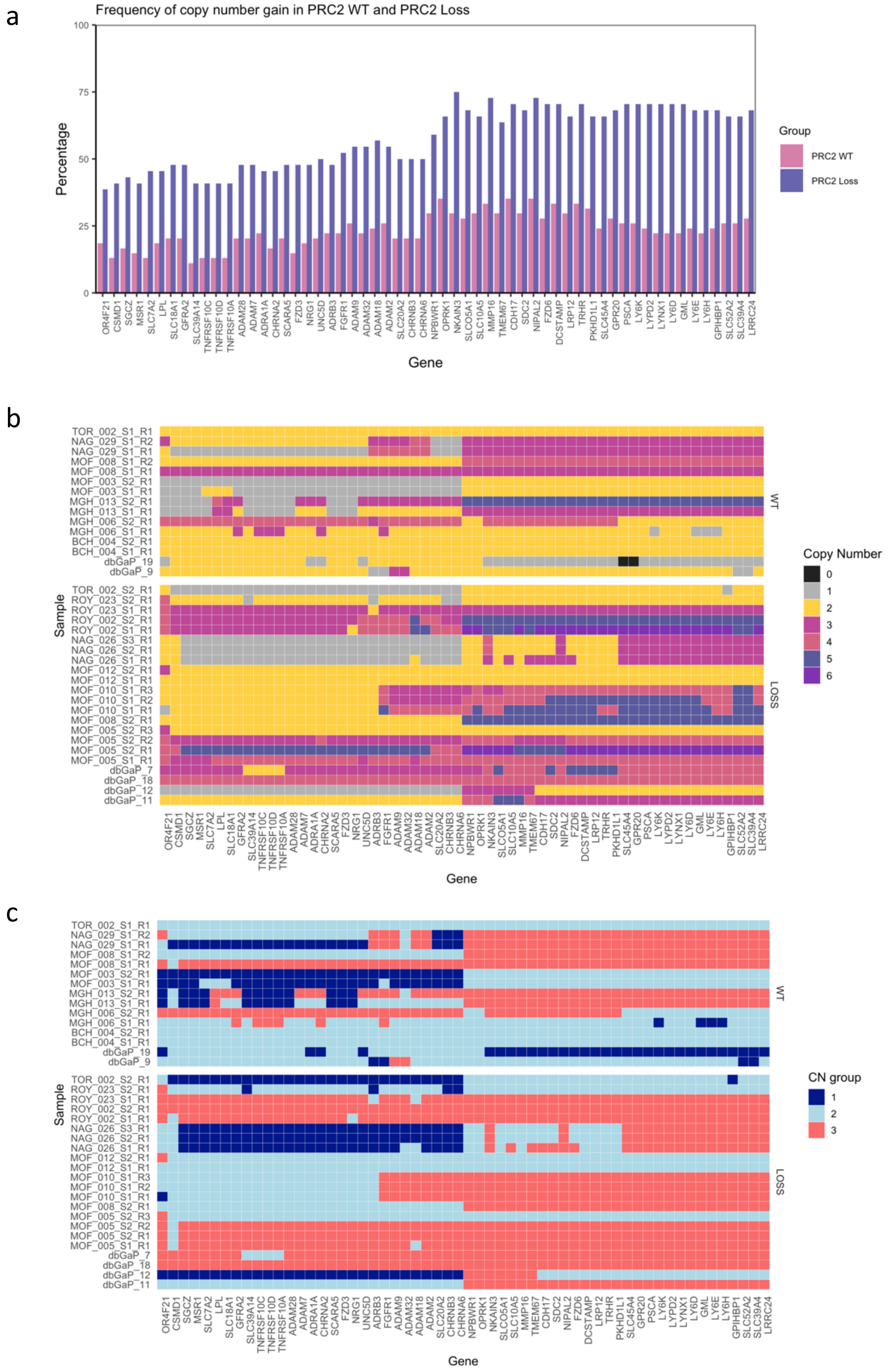


**Supplementary Figure 6.**

**Copy number alterations of chromosome 8 stratified by PRC2 status in MPNST.**

**a**.Bar plots showing the percentage of tumours with copy number gain (CN ≥ 3) for chromosome 8 surface genes, ordered by chromosomal position, in PRC2-WT (n=54 cases, pink) and PRC2-loss (n=44 cases, blue) tumours. Percentages were calculated at the case level. **b-c.** Heatmaps showing sample-level for chromosome 8 surface genes in cases with more than one tumour sample, stratified by PRC2 status with absolute copy number value (**b**), and copy number states (loss, neutral, gain) in **c.** dbGaP multiple samples per case are (dbGaP-12/dbGaP-11 , dbGaP-7/dbGaP-18, and dbGaP-9/dbGaP-19), and GeM multiple samples per case share the same 6 identifier (e.g MOF_005).

a

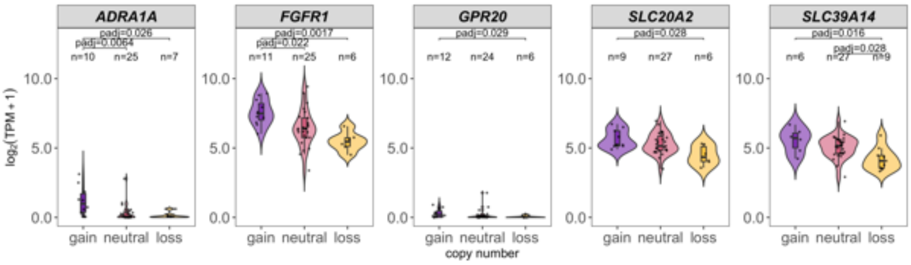
ADRA1A
FGFR1
GPR20
SLC20A2
SLC39A14
log2(TPM + 1)
gain neutral loss
copy number

b

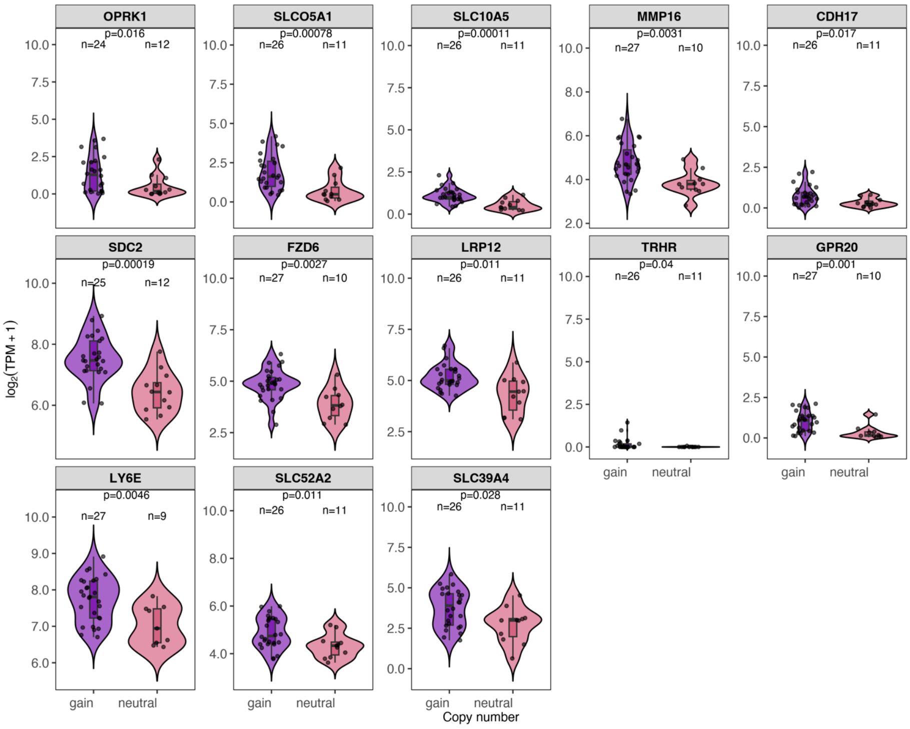
OPRK1
p=0.016
SLCO5A1
p=0.00078
SLC10A5
p=0.00011
MMP16
p=0.0031
CDH17
p=0.017
SDC2
p=0.00019
FZD6
p=0.0027
LRP12
p=0.011
TRHR
p=0.04
GPR20
p=0.001
LY6E
p=0.0046
SLC52A2
p=0.011
SLC39A4
p=0.028
log2(TPM + 1)
gain neutral
Copy number

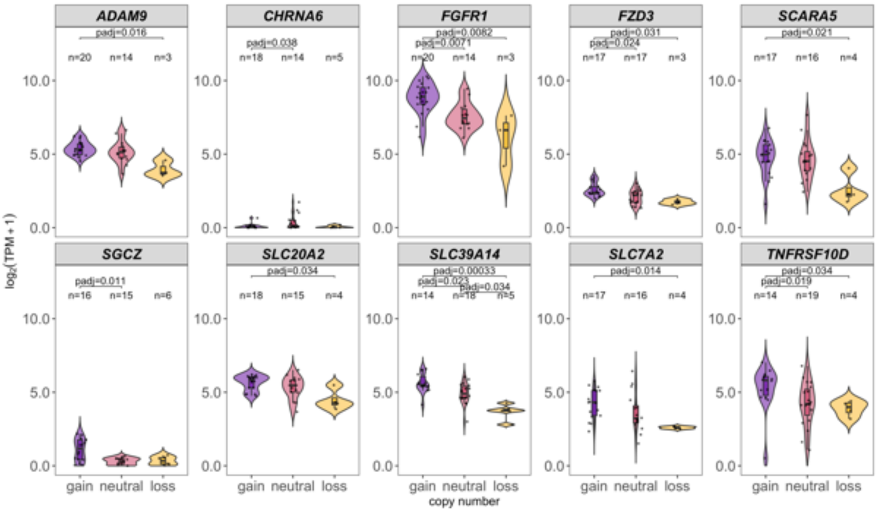
ADAM9
CHRNA6
FGFR1
FZD3
SCARA5
SGCZ
SLC20A2
SLC39A14
SLC7A2
TNFRSF10D
log2(TPM + 1)
gain neutral loss
copy number

c

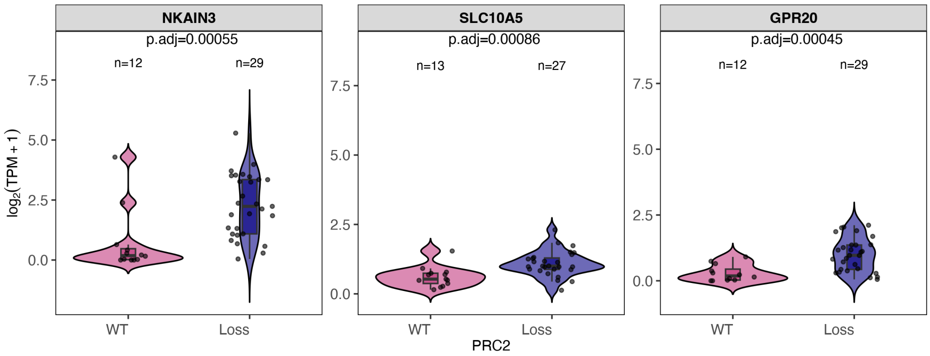


**Supplementary Figure 7.**
**Copy number alterations and expression patterns of chromosome 8 cell surface genes stratified by PRC2 status in MPNST.**

**a–c.** Violin plots showing mRNA expression of chromosome 8 surface genes in MPNST tumours. For cases with multiple tumour samples, expression values were averaged to obtain a single mean value per case prior to statistical testing. **a**. Expression in PRC2-WT tumours stratified by copy number state: gain (purple), neutral (pink), and loss (yellow). Differences in expression across copy number states were assessed using the Kruskal-Wallis test. For genes with unadjusted $p<0.05$, pairwise comparisons were performed using Dunn's test, and Holm-adjusted p-values are shown on the plot. **b**. Expression in PRC2-loss tumours. Upper panel shows genes analysed using two copy number categories (gain vs neutral), where loss was absent or represented by a single case; statistical significance was assessed using the Wilcoxon rank-sum test, and genes with $p<0.05$ are shown. Lower panel shows genes represented across all three copy number states (gain, neutral, and loss), statistical test and coloured as in panel (**a**). **c**. Comparison of surface genes with copy number gain between PRC2-WT (pink) and PRC2-loss (blue) tumours. Statistical significance was assessed using the Wilcoxon rank-sum test, and adjusted p-values (FDR) are shown. For all panels, the y-axis represents $log_2$(TPM+1), and case numbers per group are indicated.

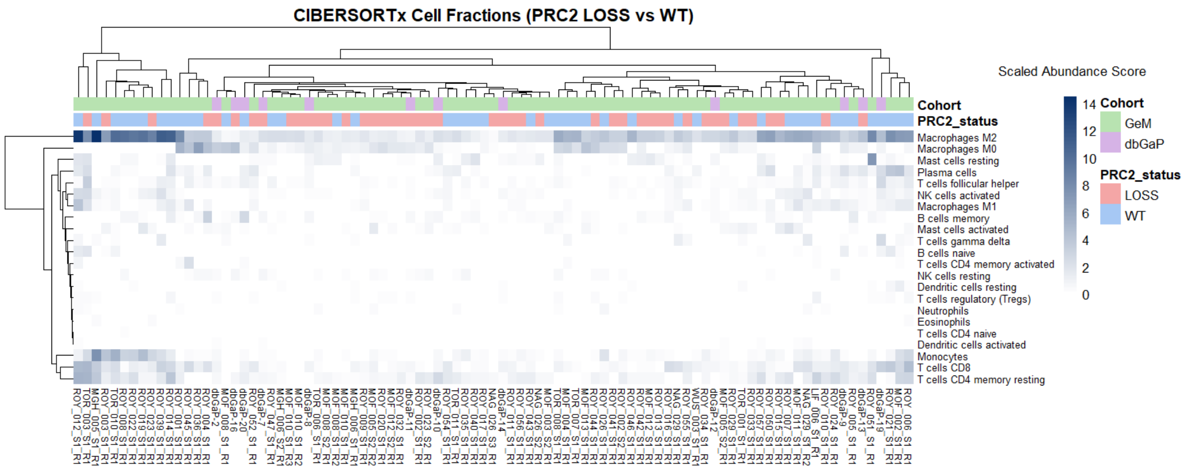


**Supplementary Figure 8.**

**CIBERSORTx immune cell subset heatmap with LM22 immune-cell signature matrix.**

Composition of immune cell subsets inferred from bulk RNA-sequencing data using CIBERSORTx with the in-built LM22 immune-cell gene signature matrix. The heatmap displays inferred scaled abundance of 22 immune cell subsets across individual samples. Samples are annotated by PRC2 status (PRC2 WT and PRC2 Loss) and data cohort (GEM and dbGAP). Hierarchical clustering was performed using complete linkage.
