## Supplementary Tables Description for "Tumour neoantigen repertoire computational predictions in malignant peripheral nerve sheath tumours define potential targets for immunotherapy"

**Supplementary Table 1.** Samples excluded from neoantigen analysis

**Supplementary Table 2.** pVACseq-predicted neoantigens with copy number data in MPNST cohorts

**Supplementary Table 3.** pVACfuse-predicted neoantigens in MPNST cohorts in MPNST cohorts

**Supplementary Table 4.** Fusion genes excluded from neoantigen prediction

**Supplementary Table 5.** Antigen-processing gene expression and Immune expression statistical analysis

**Supplementary Table 6. Chromosome 8 surface antigen expression statistical analysis**
